# A multi-modal proteomics strategy for characterizing posttranslational modifications of tumor suppressor p53 reveals many sites but few modified forms

**DOI:** 10.1101/455527

**Authors:** Caroline J. DeHart, Luca Fornelli, Lissa C. Anderson, Ryan T. Fellers, Dan Lu, Christopher L. Hendrickson, Galit Lahav, Jeremy Gunawardena, Neil L. Kelleher

**Affiliations:** National Resource for Translational and Developmental Proteomics, Northwestern University, Evanston, IL, 60208, USA; Ion Cyclotron Resonance Program, National High Magnetic Field Laboratory, Tallahassee, FL, 32310, USA; Department of Systems Biology, Harvard Medical School, Boston, MA, 02115, USA; Department of Chemistry and Biochemistry, Florida State University, Tallahassee, FL, 32310, USA; Departments of Chemistry, Molecular Biosciences and the Division of Hematology-Oncology, Northwestern University, Evanston, IL, 60208, USA

**Keywords:** p53, top-down proteomics, post-translational modifications, modforms, 21 T FT-ICR

## Abstract

Post-translational modifications (PTMs) are found on most proteins, particularly on “hub” proteins like the tumor suppressor p53, which has over 100 possible PTM sites. Substantial crosstalk between PTM sites underlies the ability of such proteins to integrate diverse signals and coordinate downstream responses. However, disentangling the combinatorial explosion in global PTM patterns across an entire protein (“modforms”) has been challenging, as conventional peptide-based mass spectrometry strategies (so-called “bottom-up” MS) destroy such global correlations. Alternatively, direct analysis of intact and modified proteins using “top-down” MS retains global information. Here, we applied both strategies to recombinant p53 phosphorylated *in vitro* with Chk1 kinase, which exhibited 41 modified sites by bottom-up MS, but no more than 8 modified sites per molecule detected by top-down MS. This observation that many low-abundance modifications comprise relatively few modforms above a 1% threshold indicates that endogenous p53 PTM complexity may be more definable than previously thought.

## Introduction

The development of nucleic acid sequencing technologies has domesticated the genome and revealed the unexpected richness of RNA biology. Proteins, however, remain central players in most cellular processes as the vast complexity of the proteome becomes better appreciated through the application of emerging technologies [1]. Aside from somatic mutations in coding sequences and alternative splicing, most proteins are also post-translationally modified, often on multiple sites. This is particularly the case for “hub” proteins of key cellular pathways, which integrate diverse input signals and help orchestrate multiple downstream responses.

The tumor suppressor p53 provides a potent example of proteomic complexity. p53 is a master cellular regulator and frequently mutated in human cancers [2-4]. It integrates input signals from myriad stressors, including genotoxic damage, oncogenesis, hypoxia, and viral infection [5, 6] into highly orchestrated downstream responses, such as apoptosis, mitosis, and metabolic changes [7-9]. It does so in a highly dynamic [10, 11] and stressor- and cell-type-specific manner [12]. The activity and stability of p53 is associated with hundreds of individual post-translational modifications (PTMs) reported to date [13-15], which are known to be regulated by numerous enzymes [13, 14] within stressed and unstressed cells. These PTMs have been identified on over 100 sites within the N-terminal transactivation domains (AA 1-92), central DNA-binding domain (AA 102-292), and C-terminal tetramerization and basic domains (AA 292-393) of the protein (**Figure 1A**) [3, 13, 14].

**Figure 1.**
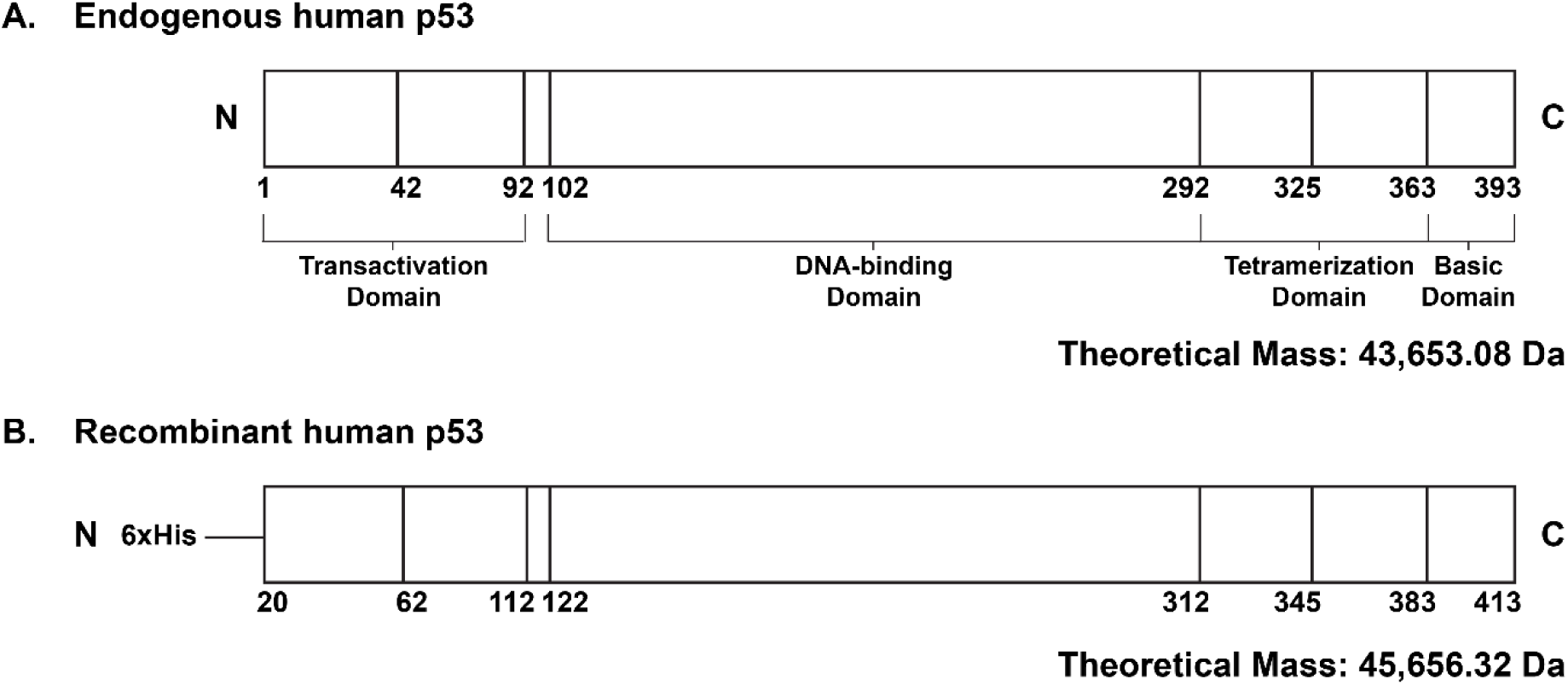
Sequence of endogenous and recombinant human p53. **A.** Endogenous human p53 sequence, with functional domains and theoretical monoisotopic mass indicated. **B.** Recombinant human p53 (rp53) sequence, comprising the full-length endogenous human p53 sequence with an N-terminal 20-residue 6xHis tag and linker sequence, with corresponding changes in residue numbers and theoretical monoisotopic mass indicated.

For p53 and similar proteins, it is likely that extensive crosstalk exists between PTM sites on different domains to integrate upstream signals and downstream responses. Such PTM crosstalk offers a way to encode information about context and function in the pattern of modifications across the whole protein, as suggested by the concept of combinatorial “PTM codes”, of which the histone code is best known [16]. Crosstalk between modified sites could arise through protein folding, which can bring widely separated parts of a peptide sequence together in space, as well as through allosteric conformational changes that could alter binding partners [17]. If such crosstalk underlies information encoding and integrative function, then it becomes essential to know the patterns of modification across the entire protein, as well as the stoichiometric abundance and distribution of these patterns within a given biological context. This presents a formidable challenge, especially for extensively modified proteins such as p53, which could theoretically exhibit over 10^30^ modified forms. While these cannot all exist, given the limitation of protein copy number in cells or entire organisms [1], the sheer scale of combinatorial PTMs highlights the challenge of determining which modified p53 forms are actually present within a given biological context.

The term “proteoform” covers all changes of a given protein molecule arising after transcription, including alternative spliceforms, chemical adducts, proteolytic cleavages, and PTMs [1, 18]. In the context of information encoding by PTMs, it is more helpful to focus on “modforms” [19], or those proteoforms arising from reversible enzymatic modification and de-modification, such as phosphorylation, acetylation and ubiquitinylation. The number of modforms that a protein can exhibit grows exponentially with the number of modification sites. An individual protein molecule will exhibit one modform, while a population of protein molecules will exhibit a distribution of modforms in varying proportions. This “modform distribution” offers the most complete information about global PTM crosstalk and a protein's capability to encode information.

Post-translational modifications of p53 are typically analyzed by PTM-specific antibodies, which provide high sensitivity but can only identify a few modifications. Traditional proteomic approaches employing proteolytic digestion and analysis of the resulting peptides offer another approach for PTM detection and are commonly referred to as “bottom-up” (BU) mass spectrometry (MS). This latter strategy excels at high-throughput, high-confidence PTM characterization of p53 populations, most notably in a pair of recent studies which performed comprehensive PTM cataloging of p53 isolated from normal human fibroblasts and identified 230 PTMs across 101 amino acid residues of p53 [15, 20]. However, proteolytic digestion severs the linkage of combinatorial PTM patterns, losing information on PTM crosstalk between functional domains and making accurate determination of complex modform distributions impossible [21].

An alternative strategy, analysis of intact modforms by “top-down” (TD) MS [22], offers a complementary approach. By keeping the complete protein primary structure intact, the PTM landscape of a proteoform can be characterized in precise molecular detail, and correlations between widely separated modification sites can be maintained [23-25]. In recent work, we showed that TD MS offers a significant advantage for estimating modform distributions [19, 21], especially when used in combination with BU MS and other “middle-down” (MD) MS approaches, which employ partial enzymatic digestion to provide ~5-25 kDa protein fragments [26-28]. However, analysis of intact endogenous human p53 modforms by TD MS presents a formidable technical challenge, due to both the molecular weight (43.6 kDa) and the sheer number of potential modforms. A method by which the number and identity of PTMs on the most abundant p53 modforms could be characterized within a specific biological context would therefore be immensely valuable to the fields of p53 biology and translational proteomics.

To explore the capabilities of TD MS for p53 modform characterization, we focused on recombinant human p53 (hereafter, “rp53”), phosphorylated *in vitro* by the kinase Chk1, which is known to target p53 *in vivo* [29-32]. Despite showing far less PTM complexity than endogenous p53, this representative model reflects the patterns of modification produced by typical regulatory enzymes. Although previous studies of Chk1 action on p53 have shown only 8 phosphorylation sites, we mapped a total of 41 phosphorylation sites to rp53 by combined BU and MD MS. However, TD MS revealed no more than 8 phosphorylation sites on each intact molecule, within the limits of detection. Moreover, as rp53 was digested into smaller peptides, the number of abundant phosphorylation sites observed on each peptide was always vastly lower than the theoretical maximum. Despite the potential complexity of 2^41^, or over 2 x 10^12^, modforms, the actual observed complexity is far less when viewed by TD and MD MS. In other words, we found many modification sites but few modforms. As we show here, TD MS offers a new approach to quantifying global patterns of PTMs and suggests that the actual PTM complexity generated and maintained by cells may be more definable than previously thought.

## Results

### Method optimization for analysis of intact p53 modforms

To the best of our knowledge, a method for the separation of intact p53 modform populations into isobaric peaks by high-performance liquid chromatography and subsequent characterization by tandem mass spectrometry (LC-MS/MS) has yet to be established. Therefore, in order to perform this critical aspect of our multimodal proteomics strategy, we first needed to determine which sample, chromatographic, instrument, and data analysis parameters would be required for rapid and high-confidence identification of intact p53 protein. To develop an LC-MS/MS method, we isolated recombinant full-length human p53 (rp53) (**Figure 1B**) from an *E. coli*-based system [33] by cobalt affinity chromatography (**Figure S1A**) and introduced it into two different Fourier-transform (FT) mass spectrometers for analysis by LC-MS/MS.

We first obtained a successful medium-resolution intact mass (MS^1^) spectrum of purified rp53 on an Orbitrap Fusion Lumos mass spectrometer, utilizing a short transient acquisition of the FT signal that enhances signals of charge states produced by electrospray ionization (ESI) using the lowest possible resolving power (7,500 at 200 *m/z*) (**Figure S2A**) [34]. After performing charge state deconvolution using the ReSpect™ algorithm, the resulting average mass of 45,683.75 Da was within 1.6 Da of that calculated from *in silico* translation of the rp53 plasmid sequence [33]. We confirmed our identification of rp53 by targeting selected charge states for fragmentation by collision-induced dissociation (CID), resulting in the generation of protein fragment ions (MS^2^). The monoisotopic masses of these fragment ions could then be calculated using the Xtract algorithm and compared against the rp53 protein sequence using ProSight Lite data analysis software [35, 36], with high-confidence matches determined by probability-based scoring metrics (e.g. p-score) (**Figure S3A**).

As we believed that a high-resolution MS^1^ spectrum would be required to accurately identify and characterize multiple PTMs on intact p53, we next performed top-down LC-MS/MS analysis of intact rp53 on the 21 Tesla (T) Fourier Transform (FT) – Ion Cyclotron Resonance (ICR) mass spectrometer at the National High Magnetic Field Laboratory (Tallahassee, FL), which holds the current world record for FT resolving power [37]. By doing so, we successfully obtained a high-resolution MS^1^ spectrum of intact rp53 (150,000 at 400 *m/z*) on a chromatographic timescale (**Figure S2B**), with charge states isotopically resolved to the baseline (**Figure S2B**, **inset**), a feat not yet achievable on any other MS platform. By performing charge state deconvolution using the Xtract algorithm, we calculated the monoisotopic mass of intact rp53 to be 45,656.45 Da, only 0.13 Da away from the theoretical mass. We obtained additional improvements in rp53 sequence coverage by employing electron-transfer dissociation (ETD) as the MS^2^ fragmentation method (**Figure S3B**), thus proving this strategy capable of rapid, high-confidence rp53 modform characterization by TD LC-MS/MS.

### Characterization of representative p53 modforms

With a workflow for accurate identification and characterization of intact rp53 by TD MS, we next analyzed a population of rp53 modforms created *in vitro* by a single enzyme. We selected purified human Chk1 kinase for this purpose because this enzyme had been previously described to phosphorylate human p53 *in vitro* and/or *in vivo* at 8 residues (Ser 15, Thr 18, Ser 20, Ser 37, Ser 313, Ser 314, Ser 378, Thr 387) [29-32], and could therefore provide up to 256 potential rp53 modforms from a single enzymatic reaction. Our rationale was to visualize the number of phosphorylated sites present on the most abundant rp53 modforms by TD, while employing complementary BU analysis to confirm the number and locations of all phosphorylated rp53 residues. For the TD approach, we subjected replicate preparations of Chk1-phosphorylated rp53 (phospho-rp53) (**Figure S1B**), in addition to kinase-free rp53 controls prepared in parallel, to LC-MS/MS analysis on an Orbitrap Fusion Lumos mass spectrometer, employing the LC and “medium-resolution” MS parameters previously optimized for unmodified rp53. Upon performing charge state deconvolution using the ReSpect™ algorithm, we observed that intact Chk1-treated rp53 modforms bore up to 8 phosphorylated sites (**Figure 2A, S4A, and S5**), consistent with the number predicted by previous studies. We further confirmed the number of detectable phosphorylated sites on rp53 modforms by performing high-resolution MS^1^ analysis of unmodified and Chk1-treated rp53 on the 21 T FT-ICR mass spectrometer. By isolating the 44+ charge state of phospho-rp53, we were able to maximize our ion signal and visualize peaks for each of the 8 rp53 phospho-modforms at near-baseline isotopic resolution (**Figure S4B**). However, due to the size and signal complexity of the target species, MS^2^ fragmentation of intact phospho-rp53 modforms could not be successfully obtained by LC-MS/MS on either instrument platform.

**Figure 2.**
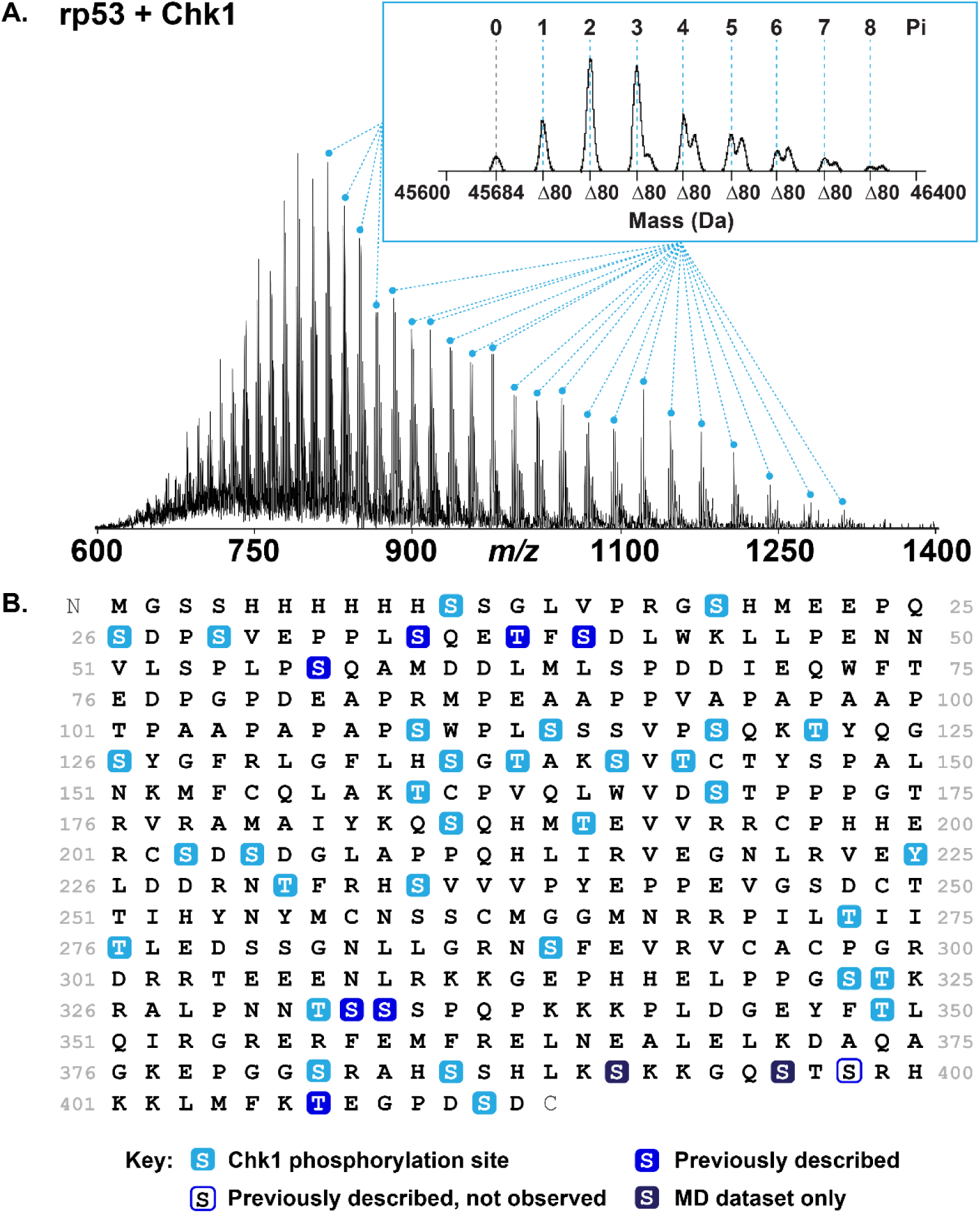
Initial characterization of representative p53 modforms. **A.** MS^1^ spectrum of *in vitro* phosphorylated recombinant human p53 (rp53) obtained by TD LC-MS/MS on an Orbitrap Fusion Lumos mass spectrometer using an FT RP of 7,500 at 200 *m/z* and a custom short-transient method. **Inset:** Average masses of the 8 rp53 phospho-modforms deconvolved using the ReSpect™ algorithm in BioPharma Finder. **B.** Phosphorylation sites mapped by BU and MD LC-MS/MS on an Orbitrap Fusion Lumos mass spectrometer.

To localize and assign phosphorylated sites within phospho-rp53 modforms, we subjected parallel preparations of Chk1-phosphorylated and kinase-free rp53 to concurrent trypsin and LysC digestion, followed by high-resolution BU LC-MS/MS analysis on an Orbitrap Fusion Lumos mass spectrometer. We then used Mascot software to search the resulting peptide and accompanying fragment ion monoisotopic masses against the rp53 sequence, as well as the UniProt *E. coli* database to rule out any false positive matches from contaminating proteins, using a series of increasingly strict parameters (detailed in *Experimental Methods*). We also performed replicate searches using the MS Amanda [38] and Target Decoy PSM Validator nodes in Proteome Discoverer, as this workflow provided orthogonal probability-based scoring metrics and allowed us to manually validate each identified phosphorylation site within the raw data. By accepting only the highest-scoring identifications from both search strategies across three independent experiments, we identified a total of 39 phosphorylated residues on rp53 resulting from *in vitro* phosphorylation by Chk1 (**Figure 2B, Table S1**). This striking increase in the number of potential phosphorylation sites from the 8 predicted by the literature proportionally increased the number of potential modforms within the sample from 256 to 5 x 10^11^ and indicated that each isobaric modform peak visualized by TD might bear far higher complexity than anticipated, thus partially explaining why MS^2^ based characterization of intact phospho-rp53 could not be achieved.

### Integration of proteomic approaches to estimate p53 modform distributions

As our TD and BU datasets indicated two highly divergent degrees of phosphorylation exhibited by phospho-rp53 modforms, we sought to find a middle ground in which we could better visualize the number and combinations of phosphorylated sites present within specific regions of the rp53 sequence. Our rationale was that partial proteolytic digestion of phospho-rp53 would provide both smaller target modforms and reduced signal complexity, thus facilitating the identification and localization of the most abundant phosphorylation sites on each intact digestion product. For this “middle-down” (MD)-MS approach, we subjected parallel preparations of Chk1-phosphorylated and kinase-free rp53 to partial digestion with LysC and analyzed the resulting peptides by high-resolution MD LC-MS/MS on an Orbitrap Fusion Lumos mass spectrometer.

For the smaller peptide fragments (≤10 kDa), we searched the resulting data against a custom UniProt-style database, comprising the rp53 sequence and all phosphorylation sites confirmed by BU, using ProSightPC 4.0 and a three-pronged search strategy (detailed in *Experimental Methods*) [39]. We then used TDCollider software to collate the results of each high-throughput search into a single output file comprising a protein sequence coverage map and accompanying MS^2^ graphical fragment maps of all rp53 digestion products identified by MD. To rule out the possibility of false identifications, both positive and negative, we subjected each identified phospho-rp53 sub-region to manual phosphorylation site validation with ProSight Lite (examples in **Figure S6D-F**). In doing so, we identified two additional rp53 phosphorylation sites (Ser 390, Ser 395) (**Figure 2B, Figure S6E, Table S1**) within a region that received poor peptide coverage by BU analysis (AA 389-398), bringing the total number of sites identified to 41 (and the total number of potential modforms to 2 x 10^12^).

For the larger (≥10 kDa) peptide fragments, we first obtained a medium-resolution MS^1^ spectrum of each species (**Figure S7, Figure S8**) to determine both the MW and the number of potential phosphorylation sites by performing charge state deconvolution using the ReSpect™ algorithm (**Table S2**). We then performed a series of targeted MS^2^ experiments employing HCD or ETD fragmentation of individual charge states, with method parameters tailored to obtain the best possible sequence coverage of each phospho-rp53 peptide. Finally, we verified that we had identified the correct peptide fragment by calculating MS^2^ fragment ion monoisotopic masses with the Xtract algorithm and comparing them against the respective regions of rp53 using ProSight Lite (**Figure S7, Figure S8**).

To integrate the phospho-rp53 modform characterization performed by TD-, BU-, and MD-MS into a constrained estimate of the total number of potential modforms present, we first calculated the relative abundance of each phosphorylated form of intact rp53 and MD digestion products spanning the entire length of the rp53 protein sequence (**Figure 3, Figure S6**). By overlaying the final relative abundance histograms against a coverage map of the rp53 sequence and all phosphorylation sites identified therein (**Figure 3, Figure S6**), we were able to localize subsets of potential modform complexity to specific regions of phosphorylated rp53. For example, of the 8 total phosphorylated sites detected on intact rp53 (**Figure 3A**), up to 8 sites could be ascribed to the N-terminal half of the protein (**Figure 3B, 3C; Figure S7**), and 5 to the C-terminal half (**Figure 3B, 3D; Figure S8**) with the number of abundant sites appearing to decrease closer to the termini (**Figure 3E, 3F; Figure S6A-C**).

**Figure 3.**
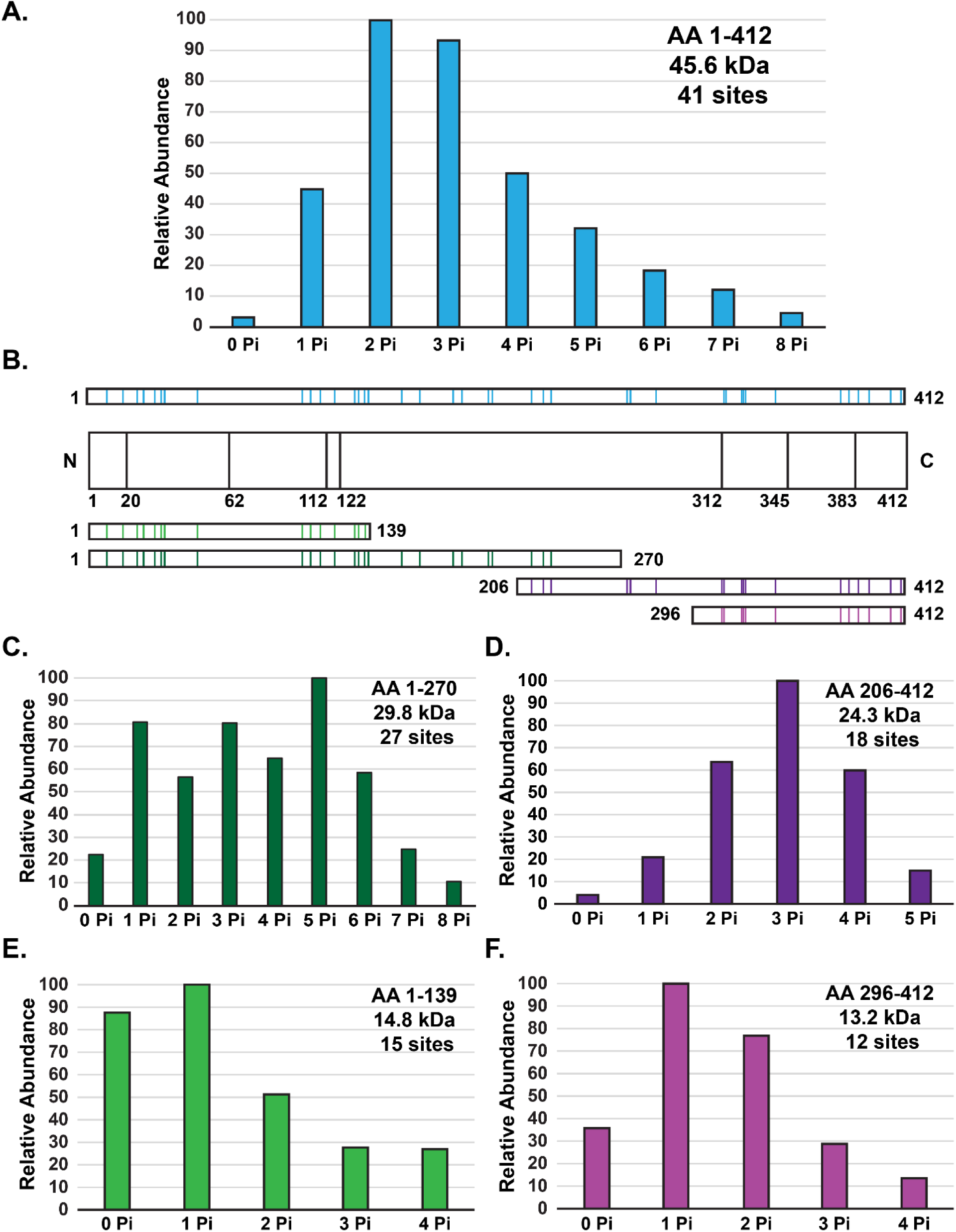
Integration of proteomic approaches to estimate p53 modform distributions. **A.** Relative abundance of all modforms of intact phospho-rp53 detected by top-down mass spectrometry. **B. Top:** Coverage map of phosphorylation sites identified by bottom-up and middle-down mass spectrometry; sites indicated with vertical lines. **Bottom:** Coverage map of the phosphorylation sites located within the largest N-terminal and C-terminal rp53 peptides identified by middle-down analysis. **C.** Relative abundance of all modforms of the N-terminal 269 amino acid residues of phospho-rp53 detected by middle-down mass spectrometry. **D.** Relative abundance of all modforms of the C-terminal 206 amino acid residues of phospho-rp53 detected by middle-down mass spectrometry. **E.** Relative abundance of all modforms of the N-terminal 138 amino acid residues of phospho-rp53 detected by middle-down mass spectrometry. **F.** Relative abundance of all modforms of the C-terminal 116 amino acid residues of phospho-rp53 detected by middle-down mass spectrometry.

By calculating the numbers of potential combinations resulting from the addition of up to 8 phosphorylated sites to intact rp53, multiplying each number by the abundance ratio calculated for the respective isobaric modform peak in BioPharma Finder, and summing the results together, we estimated a total of 68 potential phospho-rp53 modforms when considering the TD data alone (**Figure 4**, **Table S3**). By comparing the number of phosphorylation sites mapped by BU with the maximum number of sites observed within each region by MD, we could further refine our estimation of overall modform complexity. For example, the presence of 12 sites identified within AA 296-412 by BU, observed as a maximum of 4 co-occurring sites by MD, indicated the presence of 495 potential combinations (**Figure 3F**). However, only 6 sites were observed in a total of 11 combinations within that region by MD. Similarly, of the 19 sites localized to AA 1-173 by BU, with 3,874 potential combinations, only 6 sites in a total of 13 combinations were observed by MD. As we consistently observed that fewer than 50% of the phosphorylation sites predicted by BU within a specific peptide sequence were detected by MD, and that these sites were detected within a small subset of all potential combinations by MD, even for the smallest peptide fragments (**Figure S6A-C**), it is more than likely that the actual intact rp53 phospho-modform distribution occupies a similarly small subset of our BU-based estimate.

**Figure 4.**
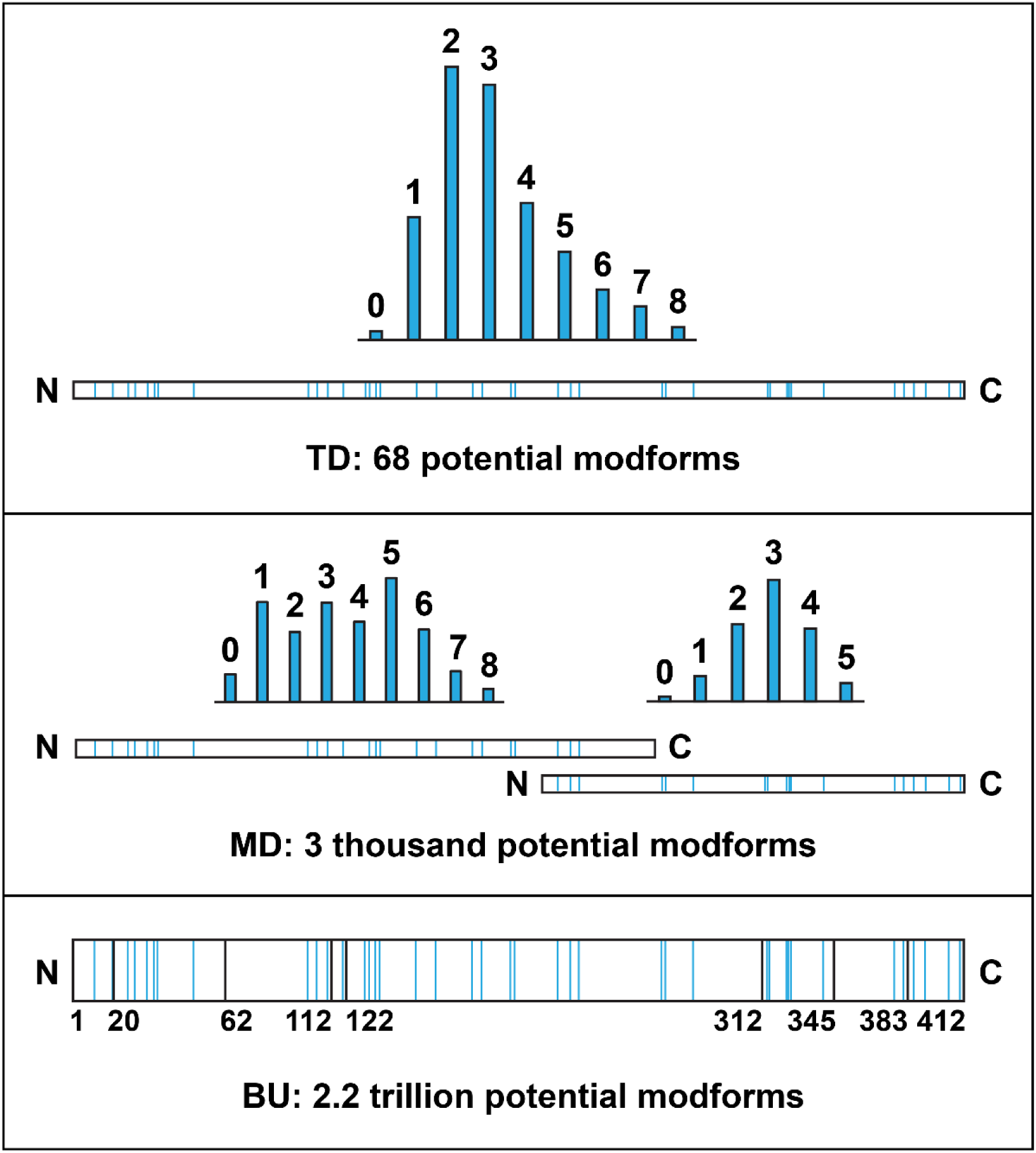
A combined strategy to characterize and constrain complex modform distributions. Using a combination of top-down (TD), middle-down (MD), and bottom-up (BU) strategies, one obtains different estimations of the maximal number of modforms present within an *in vitro* modified recombinant p53 population. One can reconcile the different estimates by applying an abundance filter and realizing that occupancies for individual PTMs can range from 100% to <0.01%; the detection of multiple sites by larger peptides creates a fractional multiplication which has PTM sites with very low occupancies contributing negligibly to the relative abundance of modforms as viewed by MS of the whole protein (which typically has a detection limit of modforms present at ^~^1% relative abundance or higher).

## Discussion

### Advancements in multimodal proteomic methodology

Single charge states of intact p53 protein are challenging to analyze by TD MS, even when abundant and unmodified. The relatively large MW of p53 (43.6 kDa for endogenous, 45.6 kDa for recombinant) is at or above the upper limit of most commercially available MS instrumentation for high-quality MS^1^ and MS^2^ fragmentation data on a timescale compatible with chromatography. This is due to the MW-dependent decrease in signal-to-noise ratio from the concomitant increase in peaks generated by electrospray ionization and chemical adducts [40], a situation exacerbated by PTMs and other sources of biological variation (e.g. mutations). Special measures, such as the “medium-resolution” method employed on the Orbitrap Fusion Lumos or the superior signal-to-noise ratio provided by the 21 T FT-ICR, must therefore be taken to obtain high mass accuracy measurements and confident identification of intact p53 protein by TD MS. Moreover, the biochemical properties of rp53 present an additional challenge, namely rapid and reproducible degradation outside of the sample parameters detailed in the Experimental Methods. Finally, the addition of PTMs at variable sites and at widely variable stoichiometries provides a significant increase in both sample and signal complexity, thus rendering the chromatographic resolution and ensuing accurate mass determination of individual rp53 modforms a formidable proposition.

After surmounting these challenges to develop a workflow by which intact rp53 could be successfully analyzed and characterized by TD LC-MS/MS on two platforms, the Fusion Lumos (**Figure S2A, S3A**) and the 21 T FT-ICR (**Figure S2B, S3B**), we were able to analyze intact and phosphorylated rp53 modforms generated by Chk1 kinase *in vitro*. While complementary PTM cataloging by high-resolution BU and MD reproducibly showed that 41 phosphorylation sites were present on Chk1-treated rp53 (**Figure 2B, Table S1**), substantially more than the literature had indicated, we were able to directly visualize by intact mass that the most abundant modforms within our sample population bore only up to 8 phosphorylated sites with a threshold of ^~^2-3% relative abundance for modform detection (**Figure 2A, Figure S4**). As viewed by whole-protein mass spectrometry, phosphorylated rp53 greatly constrained our estimate of how many rp53 modforms were present above an abundance threshold and vastly diminished the degree of potential modform complexity predicted from the BU data alone. Moreover, in establishing the parameter requirements for the analysis of high-MW, highly-modified intact proteins by LC-MS/MS, we have laid a path for the characterization of complicated modform populations from a broad range of biological contexts.

The intermediate nature of the MD approach also provides numerous benefits when attempting to characterize a complex modform population. Foremost, peptides from ^~^5-20 kDa are smaller and less extensively modified than whole modforms, thus facilitating localization of abundant modification sites by standard MS^2^ fragmentation techniques. While partial digestion products > 20 kDa may not be as accessible to MS^2^ characterization (**Figure S7, Figure S8**), due to the number of potential fragment ion channels exceeding the capacity of the mass analyzer (Orbitrap Fusion Lumos), they can still indicate the number of abundant modification sites detectable within a given region of the protein (**Figure 3**). This latter point proved especially helpful when estimating the total number of rp53 phospho-modforms, as the number of abundant modifications observed per region was always significantly less (50% or below) than the number predicted by BU (**Figure 2B, Figure 3**), even for the smallest pieces analyzed (**Figure S6**). While our final rp53 modform estimate was somewhat impacted by lack of complete MS^2^ sequence coverage within the largest phosphorylated rp53 peptides, which spanned the central 100 AA of rp53, MD analysis indicated that most of the phosphorylation sites were likely to exist closer to the N- and C-termini (**Figure 3**), from which we were able to obtain high sequence coverage by MS^2^ fragmentation and therefore complete localization of multiple phosphorylation sites.

### Biological implications on p53: many modifications, few abundant modforms

p53 is a transcription factor and nodal signal conduit protein responsive to a diverse repertoire of upstream cellular stresses (e.g. hypoxia, oncogenic signaling, DNA damage) while regulating myriad downstream effector pathways encompassing thousands of target genes [41] collectively coordinating the final cellular response (e.g. DNA damage repair, apoptosis, cell cycle arrest). An explanation for how a single protein can perform such a wide range of roles in a context specific manner has been provided by the discovery of additional PTMs present on p53. Indeed, several specific enzymatic phosphorylation or acetylation events have been shown to alter the cell fate selection by p53 [42-45]. More recently, endogenous human p53 has been shown to bear more than 300 PTMs at individual sites with unknown stoichiometries resulting from stressor-specific cellular response pathways [13-15]. This has led to a prevailing speculation of the existence of a highly complex PTM-based combinatorial logic, or “PTM barcodes” [12]. Patterns of PTMs are thought to direct the ability of p53 to activate a response to a given stressor in a cell by localizing to the nucleus and binding to a small subset of over 3500 transcriptional targets [7, 46], often by interacting with only a few protein binding partners out of potential thousands [13, 41]. Given the sheer number of possible PTM combinations (estimated at 10^30^), it is difficult to envision a feasible evolution of an equal number of ‘readers’ for every PTM combination, due to the metabolic burden this would impose on any given cell. However, questions remain: just how complex is the p53 PTM code and how many p53 modforms are there?

Pre-existing techniques such as immunoblots or even BU proteomics have not been able to reveal the true scope of PTM-based p53 regulation over decades of dedicated research. Analysis of p53 populations by modification-specific antibodies or standard BU proteomics methods only indicate which PTMs might be present; elucidation of PTM crosstalk between different regions of the protein, the modform distribution across a p53 population, or the PTM logic correlating to p53 modform function within any given biological context is far more challenging. Indeed, when comprehensive peptide-based PTM catalogs of equivalent p53 populations resulting from different stressors were compared, there were no significant differences observed in the location or identity of any PTMs, despite each population exhibiting distinct functional phenotypes (e.g. ability to activate transcription of downstream target genes) [20]. Without a method to determine which p53 modforms are expressed at stoichiometric abundance as the result of a specific cellular stressor, elucidation of the underlying PTM logic governing even a single p53-dependent response pathway will likely remain elusive. Our TD MS workflow for intact p53 analysis therefore provides a first attempt to decipher modforms directly on whole p53 molecules.

In order to reduce the complexity and competition of multi-enzyme modifications on p53, we limited our study to *in vitro* phosphorylation by a single DNA damage responsive kinase, Chk1. While we do not know if the 33 new phosphorylation sites on rp53 reported here are physiologically relevant, our collective BU, MD, and TD MS results nevertheless provide new insight into the readout of PTM-based logic. Firstly, we detected a range of individual rp53 molecules containing combined totals of 0 through 8 phosphorylation events by TD. This reinforces the hypothesis that PTMs modulate p53 activity in a graded, rheostat-like response [47], wherein the activity of p53 is more critically ascribed by the number of phosphorylation sites, rather than by their precise locations. Secondly, by constraining our estimated modform distribution to the empirical data provided by our integrated proteomic approach, we were able to narrow our estimate to 68 potential rp53 modforms, approximately 11 orders of magnitude lower than the 2 x 10^12^ potential modforms resulting from a total occupancy of all 41 phosphorylation sites (**Figure 4**). This emphasizes that the actual number of modforms present within a population is substantially less than the total number of theoretically possible combinations, thus significantly reducing the potential complexity of the regulatory PTM barcode.

The next step, which we have already began to investigate, will be to employ mathematical algorithms to compute the precise abundance of all empirically observed modforms within a given population and thereby constrain the distribution of all potential p53 modforms. This ability to constrain and quantify p53 modform distributions using an integrated MS approach will facilitate the identification of potential redundancies or linkages between modification sites, providing greater insight into the underlying logic of signal pathways and p53 PTM codes. Until TD MS technology is capable of performing complete PTM localization of complex p53 modforms, we must rely on complementary proteomic strategies to provide the most conservative estimate of p53 modform distributions and constrain the complexity of the p53 PTM landscape. While the kinase-treated samples studied herein bear dozens of low-abundance PTM sites from promiscuous activity *in vitro*, the modform distribution created *in vivo* is likely to be quite different. *In vivo*, one could expect tighter control over modification site utilization, but also that some degree of site promiscuity and biochemical “noise” is inevitable. Advancements in measurement science should aim to quantify endogenous p53 modform distributions, clarifying the spatiotemporal crosstalk among modifications on p53 [14] and PTM alterations over different temporal phases of p53 expression in response to stress (e.g. the pulsatile expression of p53 in response to ionizing radiation in MCF7 cells [10, 11]). These insights will in turn significantly advance our understanding of how these PTMs may change as a result of oncogenic p53 mutations, as well as the impact of the resulting p53 onco-modforms in human cancers.

## Acknowledgements

This work was supported by the National Institute of General Medical Sciences R01GM105375 (J.G., D.L. and G.L.), and P41GM108569, for the National Resource for Translational and Developmental Proteomics (NRTDP) based at Northwestern University (N.L.K.), as well as by the National Science Foundation Cooperative Agreement No. DMR-1157490 and the State of Florida (L.C.A. and C.L.H.). Further support was provided by the National Center for Research Resources award #S10RR027842 to the Keck Biophysics Facility at Northwestern University. The content is solely the responsibility of the authors and does not necessarily represent the official views of the National Institutes of Health.

## Author Contributions

C.J.D., L.F., and L.C.A. designed and conducted the experiments. C.J.D., L.F., L.C.A., and R.T.F. analyzed the data. C.L.H. and N.L.K. contributed analytical tools. C.J.D., L.F., D.L., G.L., J.G., and N.L.K. wrote the paper.

## Declaration of Interests

R.T.F., J.G., and N.L.K. are involved in the production of freeware and commercialized software, so therefore declare a conflict of interest.

## Experimental Methods

### Expression and isolation of recombinant human p53

A pET15-based plasmid allowing IPTG-inducible expression of intact human p53 bearing an N-terminal 6xHis tag [33] was purchased from Addgene (Plasmid # 24859), amplified in DH5α *E. coli* by liquid culture in LB medium, isolated via Qiagen midiprep kit (Qiagen, Hilden, Germany), verified to contain the correct p53 sequence by Sanger sequencing at the Northwestern University Genomics Core Facility (Evanston, IL), and transformed into BL21-DE3 competent *E. coli* (New England Biolabs, Ipswich, MA). Intact recombinant human p53 (rp53) was then expressed and isolated using a method modified from [33]. Briefly, transformed cells were amplified in 5 mL LB cultures overnight at 37 °C, diluted 1:50 into 150 mL LB cultures, amplified for 8 h at 37 °C, induced with IPTG at a final concentration of 0.5 mM, and transferred to 25 °C for overnight rp53 expression (typically 16-18 h). Cells were pelleted by centrifugation at 5 krpm for 30 min. at 4 °C, frozen in LN_2_ for 30s, and lysed into 25 mM Tris-HCl, pH 7.5, containing 500 mM NaCl, 10 mM imidazole, 1% Triton X-100, 4 μM leupeptin, 1.7 μM antipain, 1.5 μM pepstatin, and 1 mM PMSF. Lysates were incubated on ice for 20 min., followed by addition of 750 U of benzonase nuclease (Sigma-Aldrich Corp., St. Louis, MO) and MgCl_2_ to a final concentration of 1 mM. Lysates were incubated for 20 min. at 20 °C, clarified by centrifugation at 7.5 krpm for 15 min. at 4 °C, and incubated with 6 mL of Pierce HisPur cobalt affinity resin (Thermo Fisher Scientific, San Jose, CA) by batch rotation for 4 h at 4 °C.

Protein-bound resin was poured into a leveled 25 mL glass column (Bio-Rad Laboratories, Hercules, CA) by gravity flow and washed successively with 3 column volumes of ice-cold 20 mM Tris-HCl, pH 7.5, containing 250 mM NaCl, 4 μM leupeptin, 1.7 μM antipain, 1.5 μM pepstatin, 1 mM PMSF, and a step gradient of 10 mM, 25 mM, or 50 mM imidazole. Bound rp53 was eluted into 1 mL fractions with 1.5 column volumes of ice-cold 50 mM Tris, pH 7.5, containing 400 mM NaCl, 150 mM imidazole, 4 μM leupeptin, 1.7 μM antipain, 1.5 μM pepstatin, and 1 mM PMSF. Aliquots from each eluted fraction were subjected to SDS-PAGE and subsequent silver nitrate stain [48] to evaluate rp53 quantity and purity (**Figure S1A**). Those fractions containing the bulk of the purified rp53 were concentrated and exchanged into ice-cold 50 mM Tris-HCl, pH 7.5, containing 150 mM NaCl, 4 μM leupeptin, 1.7 μM antipain, 1.5 μM pepstatin, and 1 mM PMSF by centrifugation in a 5 mL 30 kDa MWCO filter (Merck Millipore, Billerica, MA) at 7.5 krpm for a total of 20 min. at 4 °C. Concentrated rp53 protein was then snap-frozen in LN_2_ and stored at −80°C for a maximum of 14 days prior to analysis.

### Phosphorylation of recombinant p53 in vitro by Chk1 kinase

Aliquots of concentrated rp53 were phosphorylated *in vitro* by recombinant Chk1 kinase (R&D Technologies, North Kingstown, RI) using a method modified from [30]. Briefly, 96 pmol aliquots of rp53 (~ 4.4 μg, as determined by BCA assay) were incubated with 10 pmol of Chk1 (544 ng) in 20 mM Tris-HCl, pH 7.5, containing 10 mM MgCl_2_, 2 mM MnCl_2_, 1 mM DTT, and 25 μM ATP for 40 min. at 30 °C. Phosphorylation was then halted by addition of DTT to a final concentration of 50 mM, followed by incubation at RT for 10 min. and for a subsequent 30 min. on ice. Typical reaction volumes were 50 μL, with kinase-free controls prepared in parallel. Aliquots from both preparations were subjected to SDS-PAGE and subsequent staining by Pro-Q Diamond phosphoprotein gel stain (Thermo Fisher Scientific) and SYPRO Ruby fluorescent protein stain (Thermo Fisher Scientific) (**Figure S1B**). Stained gel bands were visualized at a wavelength of 580 nm on a Typhoon 9400 phosphorimager (Amersham Biosciences, Little Chalfont, UK) at the Northwestern University Keck Biophysics Facility (Evanston, IL), or at 610 nm on a ChemiDoc XRS+ molecular imager (Bio-Rad Laboratories), respectively. Fluorescent gel images were analyzed and colorized using ImageJ [49].

### Sample preparation for denaturing top-down mass spectrometry

Aliquots of unmodified, phosphorylated, and kinase-free control rp53 were subjected to MeOH:CHCl_3_:H_2_O precipitation directly prior to LC-MS/MS analysis using a method modified from [50]. Briefly, proteins were precipitated in 8 volumes of methanol (Optima-grade, Thermo Fisher Scientific or LC-MS grade, Honeywell-Burdick & Jackson, Muskegon, MI), 2 volumes of ACS-grade chloroform (Sigma-Aldrich Corp.), and 6 volumes of water (Optima-grade, Themo Fisher Scientific or LC-MS grade, Honeywell-Burdick & Jackson), followed by phase separation via centrifugation at 13.2 krpm for 15 min. at RT. Pellets were washed twice in 8 volumes of ice-cold methanol with repeated centrifugation and immediately resuspended in 25 µL of ice-cold Mobile phase A (4.8 % ACN and 0.2% FA in Optima or LC-MS grade H_2_O). Samples were centrifuged for 5 min. at 13.2 krpm and 4 °C, transferred into pre-chilled vials, and immediately carried on ice for MS analysis. All samples were maintained on ice between LC-MS runs. Replicate samples were prepared at NRTDP (Evanston, IL) or shipped frozen to NHMFL (Tallahassee, FL) for on-site preparation and analysis.

### Sample preparation for middle-down mass spectrometry

Aliquots of phosphorylated and kinase-free rp53 samples previously analyzed by or prepared in parallel to those analyzed by denaturing top-down MS were transferred to LoBind tubes (Eppendorf, Hauppauge, NY) and precipitated in 6 volumes of acetone overnight at −80°C. Proteins were pelleted by centrifugation at 13.2 krpm for 10 min. at 4 °C, washed in an equivalent volume of ice-cold acetone with repeated centrifugation, and dried for 1 min. at RT. Pellets were resuspended in 20 μL 8M urea, reduced with DTT (10 mM final concentration) for 20 min. at RT, alkylated with IAA (36 mM final concentration) for 30 min. at RT in the dark, diluted to 150 μL in 0.1M NH_4_HCO_3_, pH 7.5, digested with 500 ng LysC (Promega) for 2 h at RT, and quenched with 3 μL formic acid (Thermo Fisher Scientific). The resulting peptides were subjected to concentration and desalting by C_4_ ZipTip (Merck Millipore), with activation performed in 100% MeOH, equilibration and washing performed in 0.1% LC-MS grade trifluoracetic acid (Sigma-Aldrich Corp.), and elution into 80% Optima-grade ACN and 0.1% trifluoracetic acid (all solutions prepared in Optima-grade H_2_O, Thermo Fisher Scientific). Eluted peptides from both preparations were diluted 1:5 in Mobile phase A and transferred to vials for analysis by middle-down LC-MS/MS.

### Sample preparation for bottom-up mass spectrometry

Aliquots of unmodified, phosphorylated, and kinase-free rp53 samples previously analyzed or prepared in parallel to those analyzed by top-down MS were transferred to LoBind tubes and precipitated in 6 volumes of acetone overnight at −80°C. Proteins were pelleted by centrifugation at 13.2 krpm for 10 min. at 4 °C, washed in an equivalent volume of ice-cold acetone with repeated centrifugation, and dried for 1 min. at RT. Pellets were resuspended in 20 μL 8M urea, reduced with DTT (10 mM final concentration) for 20 min. at RT, alkylated with IAA (36 mM final concentration) for 30 min. at RT in the dark, diluted to 150 μL in 0.1M NH_4_HCO_3_, pH 7.5, digested with 500 ng Trypsin-LysC (Promega, Madison, WI) overnight at 30 °C, and quenched with 3 μL formic acid (Thermo Fisher Scientific). The resulting peptides were subjected to concentration and desalting by C_18_ ZipTip (Merck Millipore), with activation performed in 90% Optima-grade ACN and 0.2% LC-MS grade formic acid, equilibration and washing performed in 0.2% LC-MS grade formic acid, and elution into 70% Optima-grade ACN and 0.2% formic acid (all solutions prepared in Optima-grade H_2_O, Thermo Fisher scientific). Eluted peptides from both preparations were dried via speedvac at RT, resuspended in 24 μL Mobile phase A, centrifuged at 13.2 krpm for 5 min. at RT to remove debris, and transferred to vials for analysis by tandem MS/MS.

### LC-MS/MS on an Orbitrap Fusion Lumos mass spectrometer

#### Top-down proteomics

Intact rp53 was analyzed by liquid chromatography on-line coupled to mass spectrometry (LC-MS/MS). Reversed-phase liquid chromatography was performed on a Dionex Ultimate 3000 chromatographic system (Thermo Fisher Scientific) using a vented tee setup described previously [34]. Mobile phase A was composed of 4.8% acetonitrile (ACN) in H_2_O with 0.2% formic acid (FA), while Mobile phase B consisted of 4.8% H_2_O in ACN and 0.2% FA. Top-down experiments were performed using in-house packed PLRP-S (5 μm, 1000 Å, Agilent, Santa Clara, CA) trap (150 µm i.d., 25 mm) and analytical columns (75 µm i.d., 250 mm length). The samples were first loaded onto the trap column at 3 µL/min. flow rate. After 10 min. of wash, proteins were separated at 300 nL/min. using the analytical column, held at 20 °C, over a 30 min. gradient of Mobile phase B from 20 to 35% followed by column wash at 95% B and re-equilibration at 10% B (total run time: 50 min.). The outlet of the analytical column was coupled to a homemade nanoelectrospray (nanoESI) source, where ionization was facilitated using a pulled capillary emitter (15 µm i.d., New Objective, Woburn, MA) connected to a high voltage union (potential set at 1.8-2 kV). The Orbitrap Fusion Lumos was operated in “intact protein mode”, with ion transfer tube temperature set at 320 °C. MS^1^ scans were recorded over a 500-2000 *m/z* window at 7,500 resolution (at 200 *m/z*) with 8 ms transients [34], averaging 20 microscans and with AGC target value of 4e5, applying a 4 V source-induced dissociation (SID). MS^2^ scans were acquired by targeting a given charge state or by selecting the top 2 most abundant precursors, isolated using the quadrupole over a 3 *m/z* window. Dynamic exclusion was set at 60 s with ±1.5 *m/z* unit tolerance. For high-energy collisional dissociation (HCD) runs, precursor ions were activated using normalized collision energy (NCE) of 23% and a stepped collision energy option (±5 steps). For collision-induced dissociation (CID) runs, precursor ions were activated using NCE = 28% and an 0.25 activation q. For high capacity electron transfer dissociation (ETD) runs [51], ion-ion interaction was set at 8 ms, with the reagent AGC target set at 7e5. All MS^2^ scans were acquired at 60,000 resolution (at 200 *m/z*) over a 400-2000 *m/z* window, averaging 4 microscans and setting the precursor AGC target value to 2e5.

#### Middle-down proteomics

Middle-down experiments were performed using in-house packed PLRP-S (5 μm, 1000 Å, Agilent) trap (150 µm i.d., 25 mm) and analytical columns (75 µm i.d., 250 mm length). The aforementioned vented tee setup was used, with the LC gradient being modified starting at 5% Mobile phase B and washing the column at 90% B. Flow rates were not changed. The Orbitrap Fusion Lumos was operated in “intact protein mode”, with ion transfer tube temperature set at 320 °C. MS^1^ scans were recorded over a 500-2000 *m/z* window using 120,000 resolution (at 200 *m/z*), averaging 20 microscans and with AGC target value of 2e5, applying a 4 V source-induced dissociation (SID). Data-dependent MS^2^ scans were acquired by selecting the top 2 most abundant precursors, isolated using the quadrupole over a 3 *m/z* window. Dynamic exclusion was set at 60 s with ±1.5 *m/z* unit tolerance. For HCD runs, precursor ions were activated using NCE=24% and a stepped collision energy option (±5% steps). For ETD runs, ion-ion interaction was set at 20 ms, with the reagent AGC target set at 5e5. For electron transfer dissociation – higher-energy collisional dissociation (EThcD) runs [52], ETD duration was reduced to 8 ms and supplemented with HCD fragmentation applying a 15 V potential to the HCD cell. All MS^2^ scans were acquired at 50,000 or 60,000 resolution (at 200 *m/z*) over a 400-2000 *m/z* window, averaging 4 microscans and setting the precursor AGC target value to 2e5.

#### Bottom-up proteomics

For bottom-up proteomics, both trap (150 µm i.d., 25 mm length) and analytical (75 µm i.d., 230 mm length) columns were in-house packed with C18 stationary phase (ProntoSIL C18AQ, 3 µm, 120 Å, Bischoff Chromatography). The aforementioned vented tee setup was used, with the LC gradient being modified starting at 2% Mobile phase B and washing the column at 85% B. Flow rates were not changed. The Orbitrap Fusion Lumos was operated as follows: survey scans (MS^1^) were collected at 120,000 resolution (at 200 *m/z*), over a 375-1800 *m/z* window. The MS^1^ AGC target was set at 4e5 charges, with max injection time of 50 ms. Precursor ions were quadrupole isolated (isolation window of 1.7 *m/z*) and subjected to two different types of tandem MS (MS^2^). Certain runs used EThcD to fragment peptide cations, with ETD followed by HCD activation of ETD products at 15 V potential. ETD was performed applying a base value of 100 ms for ion-ion interaction time (modulated using the precursor charge-dependent function), a maximum injection time of 80 ms and a cation AGC target of 2e5 charges. The reagent AGC was set at 5e5 charges. Alternatively, a decision tree [53] was used, where peptides fragmented by HCD (NCE set at 29%, cation AGC set at 1e5, maximum injection time 70 ms) could undergo a second isolation and MS^2^ fragmentation event performed via ETD if a neutral loss corresponding to the detachment of a phosphate group was detected during the HCD MS^2^ event. ETD was performed with the same parameters used for EThcD. All MS^2^ scans were recorded in the Orbitrap at 15,000 resolution (at 200 *m/z*) over a 110-2000 *m/z* window. All data-dependent scans were queued using a “total cycle time” logic limiting the duty cycle to 2 s. The dynamic exclusion was applied, with 45 s duration and 10 ppm precursor tolerance.

### LC-MS/MS on a 21 Tesla FT-ICR mass spectrometer

#### Top-down proteomics

For analysis of intact unmodified, phosphorylated, and kinase-free rp53 by denaturing top-down MS on the 21 T FT-ICR mass spectrometer at NHMFL [37, 54], reversed-phase liquid chromatography was performed on an ACQUITY M-class chromatographic system (Waters, Milford, MA) using the vented tee setup described above. Mobile phase A was composed of 4.8% acetonitrile (ACN) in H_2_O with 0.3% formic acid (FA), while Mobile phase B consisted of 4.8% H_2_O in 47.5% ACN, 47.5% IPA, and 0.3% FA. Top-down experiments were performed using in-house packed PLRP-S (5 μm, 1000 Å, Agilent) trap (150 µm i.d., 20 mm) and analytical columns (75 µm i.d., 250 mm length). The samples were first loaded onto the trap column at 2.5 µL/min. flow rate, and after 10 min. of wash proteins were separated at 300 nL/min. using the analytical column, maintained at room temperature, over gradient of Mobile phase B from 5 to 50% in 2.5 min., 50% to 60% in 22.5 min., 60% to 90% in 25 min., and 90% to 5% in 5 min. (total run time: 55 min.). The outlet of the column was coupled to a homemade nanoESI source, where ionization was reached using a pulled capillary emitter (15 µm i.d., New Objective) connected to a high voltage union (potential set at 1.8-2 kV). The 21 T FT-ICR was operated in “proteomics mode”, with ion transfer tube temperature set at 320 °C. Broadband MS^1^ scans were recorded over a 700-2000 *m/z* window using 150,000-300,000 resolution (at 400 *m/z*), averaging 2-4 microscans and with AGC target value of 3e6, applying a 15 V source-induced dissociation (SID). Isolated MS^1^ scans were recorded over an *m/z* window of 950-1250 (centered around the rp53 44+ charge state), using 300,000 resolution (at 400 *m/z*), averaging 4 microscans and with AGC target value of 2.25e6, applying a 15 V source-induced dissociation (SID). MS^2^ scans were acquired by targeting a given charge state or selecting the top-2 most abundant precursors within a 15 *m/z* isolation window. Dynamic exclusion was set at 120 s with ±3 *m/z* unit tolerance. For CID runs, precursor ions were activated using NCE=35% and an 0.25 activation q. For front-end ETD (fETD) runs [55], ion-ion interaction time was set at 7.5-10 ms, with 10-15 multiple fills within the multipole storage device (MSD, [37]) and the reagent AGC target set at 4e5. All MS^2^ scans were acquired at 150,000 resolution (at 400 *m/z*) over a 300-2000 *m/z* window, averaging 2 microscans and setting the precursor AGC target value to 6e5.

### Data analysis

#### Top-down proteomics

Data acquired from intact rp53 by denaturing top-down LC-MS/MS were analyzed as follows: for the medium-resolution FTMS1 spectra of unmodified or phosphorylated rp53 acquired at NRTDP, precursor average mass was determined from the MS1 spectrum by BioPharma Finder (v3.0, Thermo Fisher Scientific) using the ReSpect™ algorithm with a target mass of 46 kDa, a maximum charge of 100+, and a minimum of 6 adjacent charges over an *m/z* range of 500-2000. For the high-resolution FTMS1 spectra of unmodified or phosphorylated rp53 acquired at NHMFL, precursor monoisotopic masses were determined by the Xtract algorithm in Xcalibur (v. 3.0.57, Thermo Fisher Scientific), with S/N ratio of 3 and a maximum allowed charge of 60+. MS2 fragment ion monoisotopic masses from both experiments were obtained by use of Xtract, with S/N ratio of 3 and a maximum allowed charge of 30+. Fragment ion masses were then compared against the rp53 sequence by ProSight Lite (v1.4, http://prosightlite.northwestern.edu/, [35, 36]), with a global 10 ppm fragment ion mass tolerance and (f)ETD (c,z), HCD (b,y), or CID (b,y) fragmentation specified as applicable.

#### Bottom-up proteomics

Data acquired from unmodified or Chk1-phosphorylated rp53 peptides by tandem MS/MS were converted from .raw to Mascot .mgf files using the DTA Generator function in COMPASS (v1.0.4.5, [56]), with precursor cleaning enabled and allowed precursor charges ranging from 2+ to 6+. The resulting files were searched against both the UniProt *Escherichia coli* database and a database comprising the rp53 sequence using Mascot (v2.5.1, Matrix Science, www.matrixscience.com) and the following parameters: tryptic peptides, up to 3 missed cleavages, precursor ion charges of 2-4+, 10 ppm precursor mass tolerance, and 0.2 Da fragment ion mass tolerance, with both fixed (Carbamidomethyl (C, + 57.02 Da)) and variable (Phosphorylation (S/T/Y, + 79.96 Da), Oxidation (M, + 15.99 Da), Deamidation (N/Q, +0.98 Da), and Gln −> pyro-Glu (N-term Q, −17.03 Da)) modifications specified. Replicate searches were also performed on each .raw file against the rp53 sequence database using the MS Amanda [38] and Target Decoy PSM Validator nodes in Proteome Discoverer 1.4.1.14 (Thermo Scientific). Parameters were as follows: tryptic peptides, up to 3 missed cleavages, peak S/N threshold of 3, precursor ion charges of 2-7+, 10 ppm precursor and fragment ion mass tolerances, strict target FDR of 0.01, relaxed target FDR of 0.05, and both fixed (Carbamidomethyl (C, + 57.02 Da)) and variable (Phosphorylation (S/T/Y, + 79.96 Da), Oxidation (M, + 15.99 Da), Deamidation (N/Q, +0.98 Da), and Gln −> pyro-Glu (N-term Q, −17.03 Da)) modifications specified. The results from both search workflows were interrogated manually for accuracy and duplication across replicate injections and preparations. Those rp53 phosphorylation sites deemed to be high-confidence identifications were then collated and incorporated into a custom search database generated by modifying the UniProt XML sequence, variation, and post-translational modification database for human p53 (http://www.uniprot.org/uniprot/P04637) to incorporate the rp53 sequence and only the mapped Chk1 phosphorylation sites with Notepad ++ (v6.9, https://notepad-plus-plus.org/).

#### Middle-down proteomics

The rp53 + Chk1 XML file was used to generate a search database in ProSightPC 4.0 (Thermo Fisher Scientific), against which the .raw files from the middle-down MS experiments could be analyzed using the High Throughput Wizard function. Each .raw file was processed using the Xtract algorithm and a custom set of parameters including a precursor mass tolerance of 0.5 *m/z*, a retention time tolerance of 2.0 min., a precursor and fragment ion S/N ratio of 3, and precursor and fragment ion maximum charges of 80+ and 30+, respectively. The resulting .puf files were searched against the rp53 + Chk1 database using a three-pronged search comprising an absolute mass search (monoisotopic masses, 2.2 Da precursor mass tolerance, 10 ppm fragment mass tolerance), followed by a biomarker search (monoisotopic masses, 10 ppm precursor and fragment ion mass tolerances, all phosphorylation sites considered), followed by the same biomarker search with ‘delta m’ mode enabled (as in [39]). Data acquired from unmodified rp53 were also searched against the rp53 + Chk1 database using an identical strategy. The search results from each individual .raw file were saved as .puf files and collated into a single .tdReport file using TDCollider v1.0 (http://proteinaceous.net/product/tdcollider/) (Proteinaceous, Inc.). In addition to filtering the results by E-value (with those above 1 x 10^-4^ excluded), each hit was manually validated for accuracy prior to inclusion in the final list of phospho-rp53 modforms. Fragment ion coverage maps of each modform were created manually by exporting each hit to ProSight Lite. Modform abundance histograms were generated by exporting the averaged MS1 spectrum of each partial digest product (most abundant region of the respective chromatographic peak) from Xcalibur QualBrowser (Thermo Fisher Scientific), performing mass deconvolution and relative modform determination in BioPharma Finder (Thermo Fisher Scientific) using the Respect™ algorithm with target mass range adjusted for each respective species, exporting the resulting average masses and relative abundances to Microsoft Excel, normalizing to the most abundant peak, and manually generating histograms from the final data. Modform distribution estimations for intact rp53 and the two largest peptides analyzed by MD MS were estimated by multiplying the percent relative abundance of each species by the potential number of modforms (2^N) for 0-8 modification sites, then summing the results (**Table S3**).

### Experimental Design and Statistical Rationale

For the analysis of intact rp53 by denaturing top-down LC-MS/MS, a total of 5 biological replicate preparations of purified protein were analyzed, with up to 6 technical replicates per preparation. This number was required due to the pronounced instability of rp53 in MS-compatible buffers, which required substantial optimization of the top-down MS workflow. For the analysis of *in vitro* phosphorylated rp53, in addition to the kinase-free controls prepared in parallel to account for any chemical adducts due to the kinase reaction, a total of 3 biological replicate preparations and up to 5 technical replicates per preparation were analyzed by top-down, middle-down, and bottom-up mass spectrometry. For the latter two methods, partial or complete proteolytic digestion was performed on samples which had previously been subjected to top-down analysis, in addition to technical replicate samples generated in parallel. This was done to determine the degree of variation in Chk1-derived recombinant p53 phosphorylation patterns between biological and technical replicates, as well as to directly identify which phosphorylation sites were present within a specific phosphorylated rp53 modform population. For analysis of the bottom-up data, only those peptides with a Mascot ion score of 30 or greater, an MSAmanda score of 150 or greater, an E-value of 1 x 10^-16^ or below, and one or more manually verified localized phosphorylation sites were included in the list of identified rp53 phosphorylation sites and the custom search database. For the analysis of the middle-down data, only those species identified with an E-value 1 x 10^-4^ or below and manually verified to have fragment ion coverage of sufficient quality to identify each rp53 phosphorylation site were considered.

## Supplemental Figures

**Figure S1.**
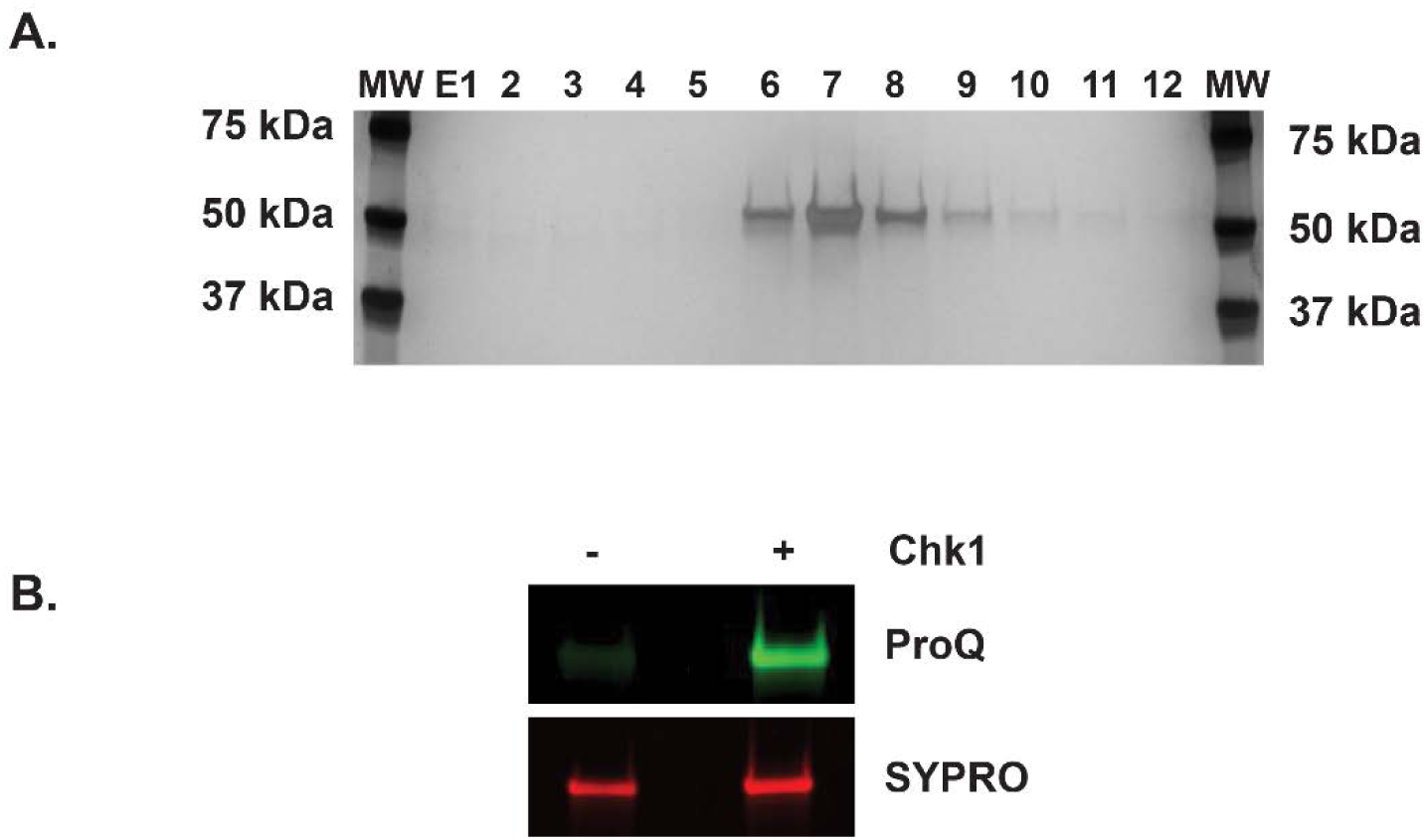
Preparing a representative p53 modform population. **A.** Silver nitrate stained gel image showing the elution profile of recombinant human p53 (rp53) purified from an *E. coli* expression system by cobalt affinity chromatography. **B.** Stained and colorized gel image showing the presence of phosphorylation on rp53 after *in vitro* modification with Chk1 kinase. ProQ (green), phosphorylated protein; SYPRO Ruby (red), total protein.

**Figure S2.**
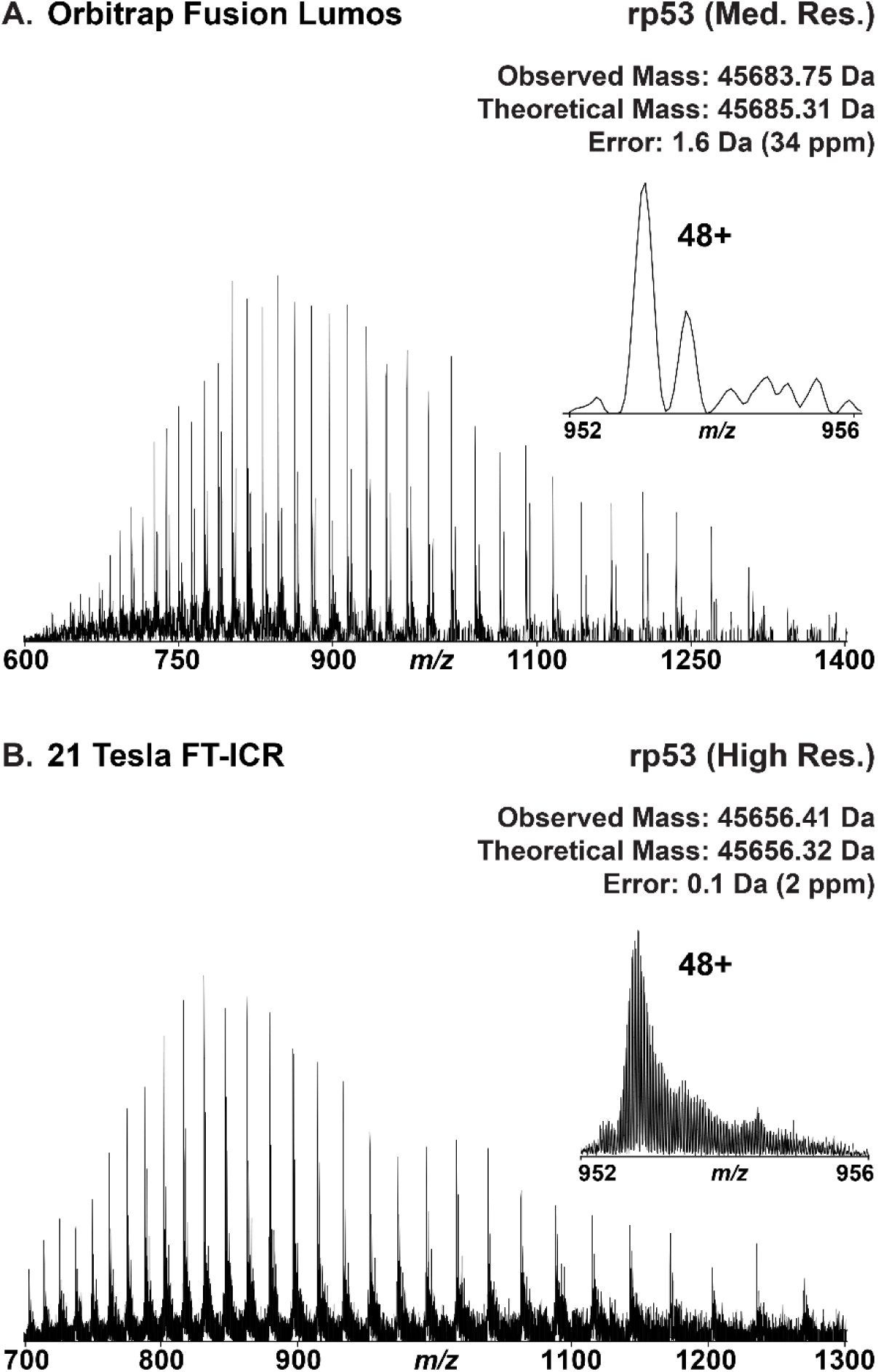
Analysis of intact recombinant human p53 by top-down LC-MS/MS. **A.** Medium resolution intact mass (MS^1^) spectrum of rp53 obtained on an Orbitrap Fusion Lumos mass spectrometer using an FT Resolving Power (RP) of 7,500 at 200 *m/z* and a custom short-transient method. **Inset:** Expansion of the 48+ charge state of rp53. **B.** High-resolution MS^1^ spectrum of rp53 obtained on a 21 Tesla FT-ICR mass spectrometer using an FT RP of 150,000 at 400 *m/z*. **Inset:** Expansion of the 48+ charge state of rp53, showing baseline isotopic resolution obtained at 21 Tesla.

**Figure S3.**
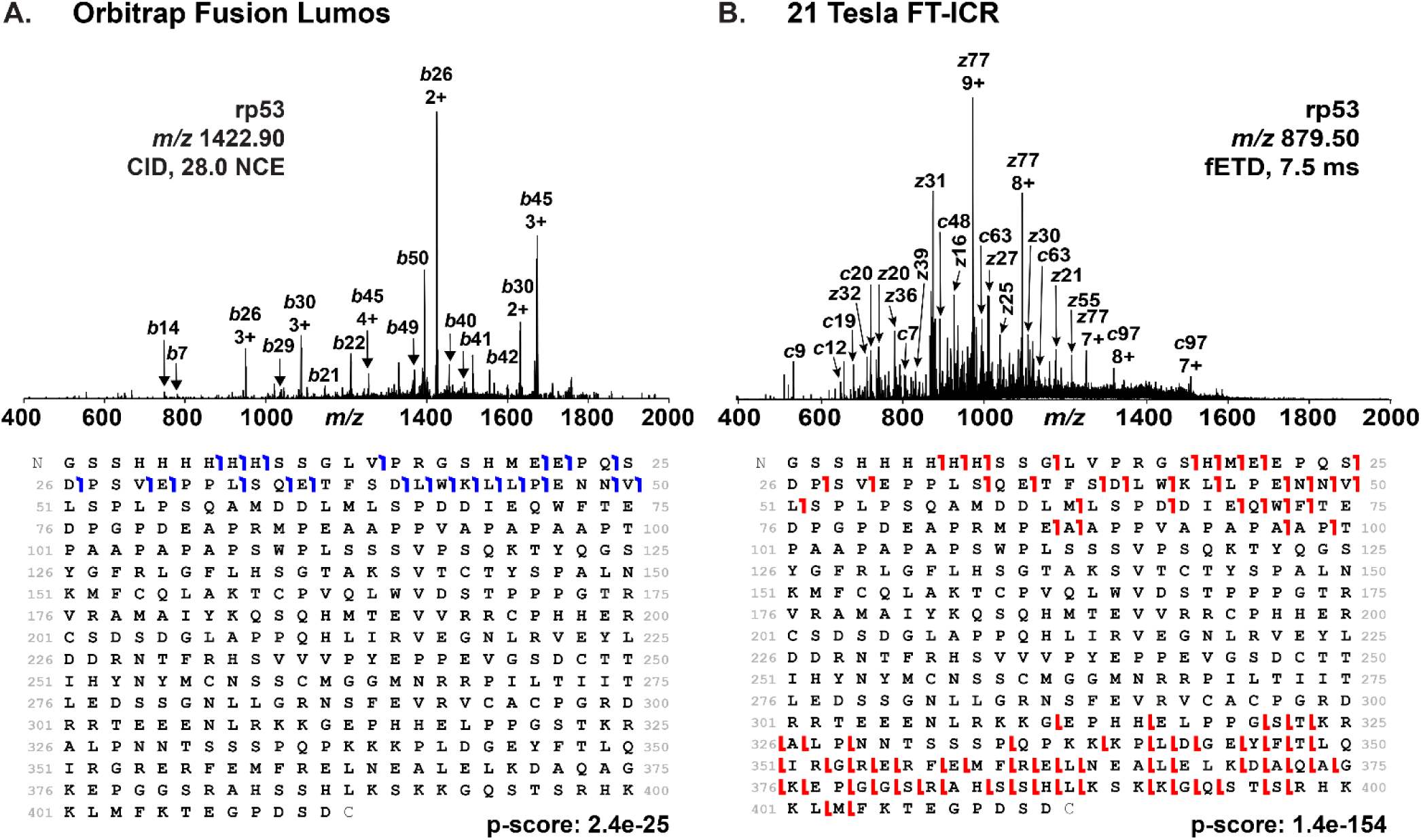
MS^2^ characterization of intact recombinant human p53 on a chromatographic timescale. **A.** Fragment ion (MS^2^) spectrum (top) and graphical fragment ion map (bottom) obtained from intact rp53 by collision-induced dissociation (CID) activation on an Orbitrap Fusion Lumos mass spectrometer. **B.** MS^2^ spectrum (top) and graphical fragment ion map (bottom) obtained from intact rp53 by front-end ETD (fETD) activation on a 21 Tesla FT-ICR mass spectrometer.

**Figure S4.**
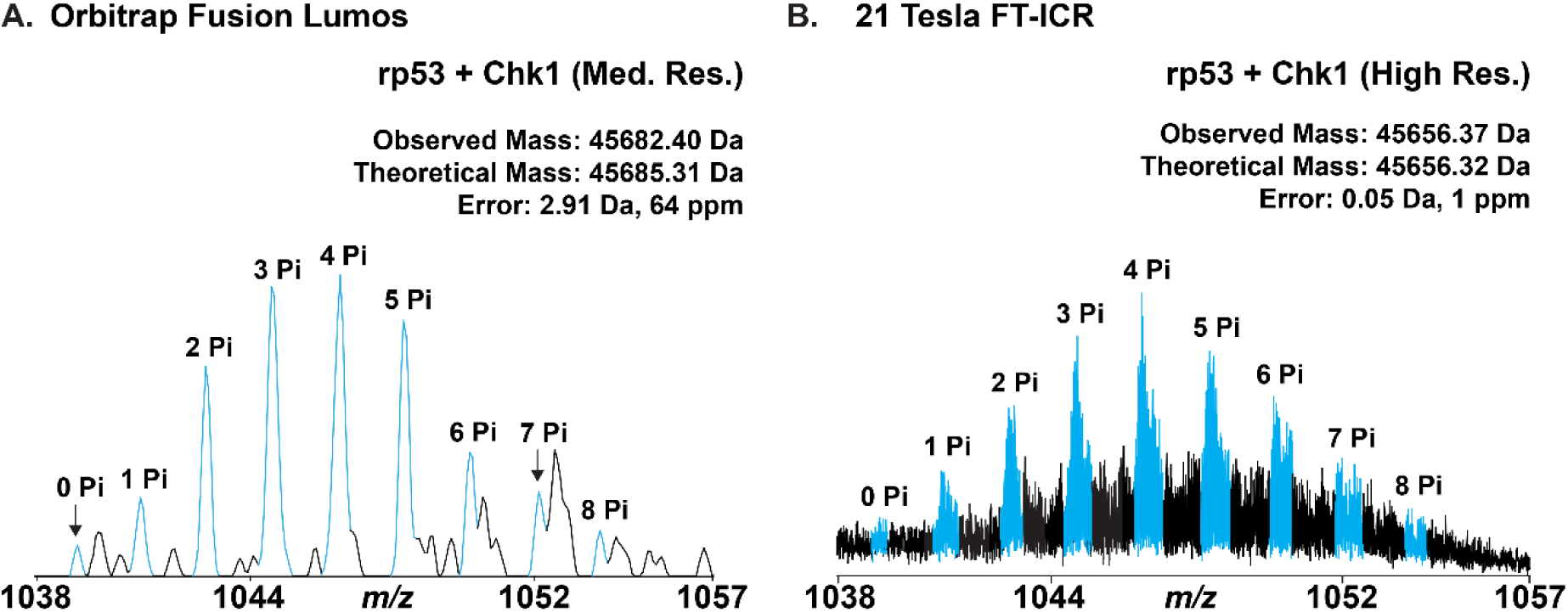
Visualization of the rp53 phospho-modform distribution by whole protein mass spectrometry. **A.** Zoomed-in view of the 44+ charge state of *in vitro* phosphorylated rp53 MS^1^ spectra obtained at medium resolution on an Orbitrap Fusion Lumos mass spectrometer. **B.** Isolated MS^1^ spectra of the 44+ charge state of *in vitro* phosphorylated rp53 obtained at high resolution on a 21 T FT-ICR mass spectrometer. For both MS^1^ spectra, peaks resulting from the addition of up to 8 phosphorylations are colored blue and labeled according to the number of modifications.

**Figure S5.**
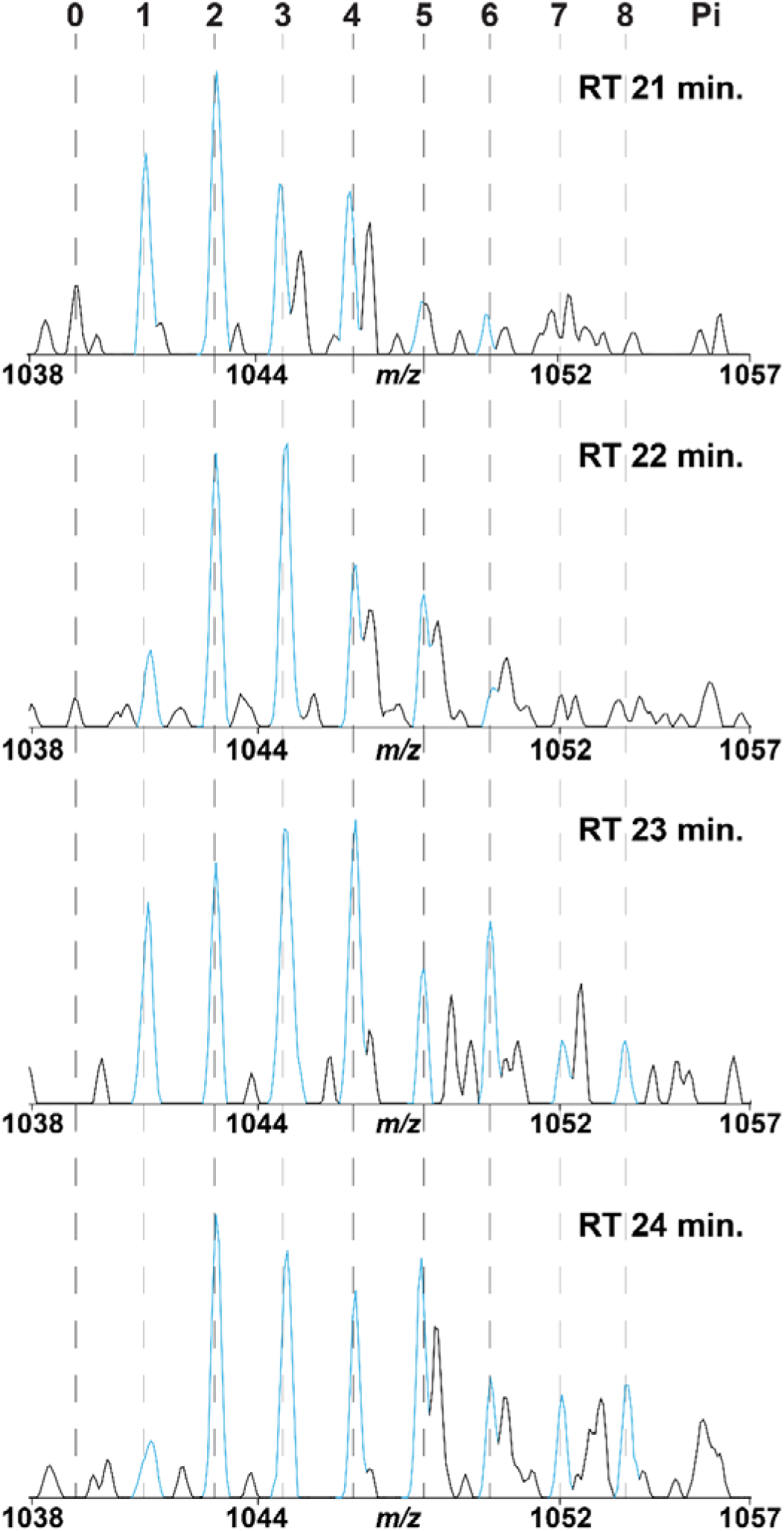
Sequential elution of intact phosphorylated p53 modforms. Zoomed-in view of the 44+ charge state of *in vitro* phosphorylated rp53 MS^1^ spectra acquired at medium resolution on an Orbitrap Fusion Lumos, sampled at sequential points along the chromatographic elution peak. RT, retention time.

**Figure S6.**
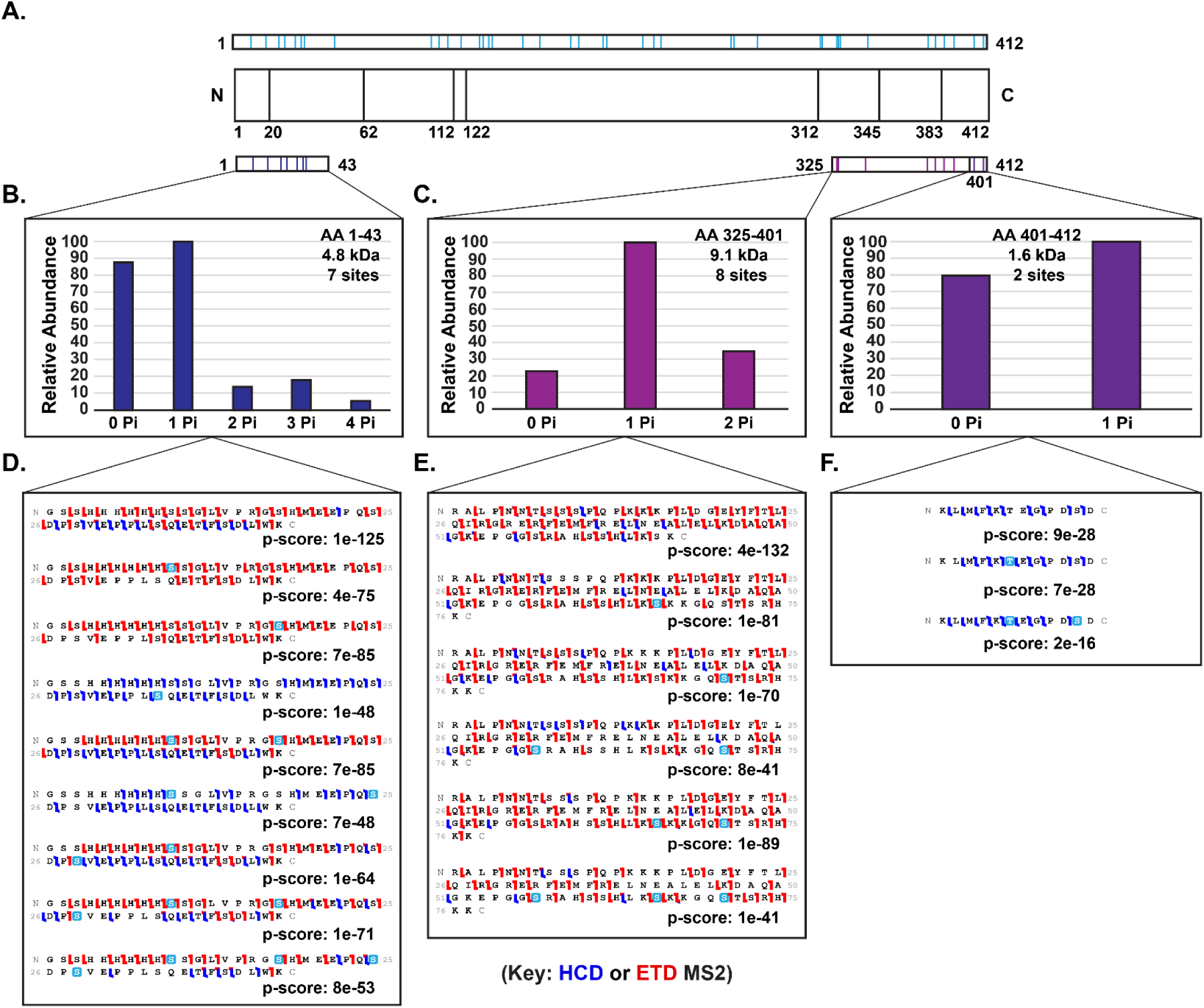
Phosphorylation sites localized to the rp53 termini by middle-down mass spectrometry. **A. Top:** Coverage map of phosphorylation sites identified by bottom-up and middle-down mass spectrometry; sites indicated with vertical lines. **Bottom:** Coverage map of the phosphorylation sites located within the smallest N-terminal and C-terminal rp53 partial digestion products identified by middle-down analysis. **B.** Relative abundance of all modforms of the N-terminal 42 amino acid residues of phospho-rp53 detected by middle-down mass spectrometry. **C.** Relative abundance of all modforms of the N-terminal 42 amino acid residues of phospho-rp53 detected by middle-down mass spectrometry. **C.** Relative abundance of all modforms of the C-terminal 87 amino acid residues of phospho-rp53 detected by middle-down mass spectrometry. **D.** Phosphorylation sites and combinations detected within the N-terminal 42 amino acid residues of phospho-rp53 by middle-down mass spectrometry. **E, F.** Phosphorylation sites and combinations detected within the C-terminal 87 amino acid residues of phospho-rp53 by middle-down mass spectrometry.

**Figure S7.**
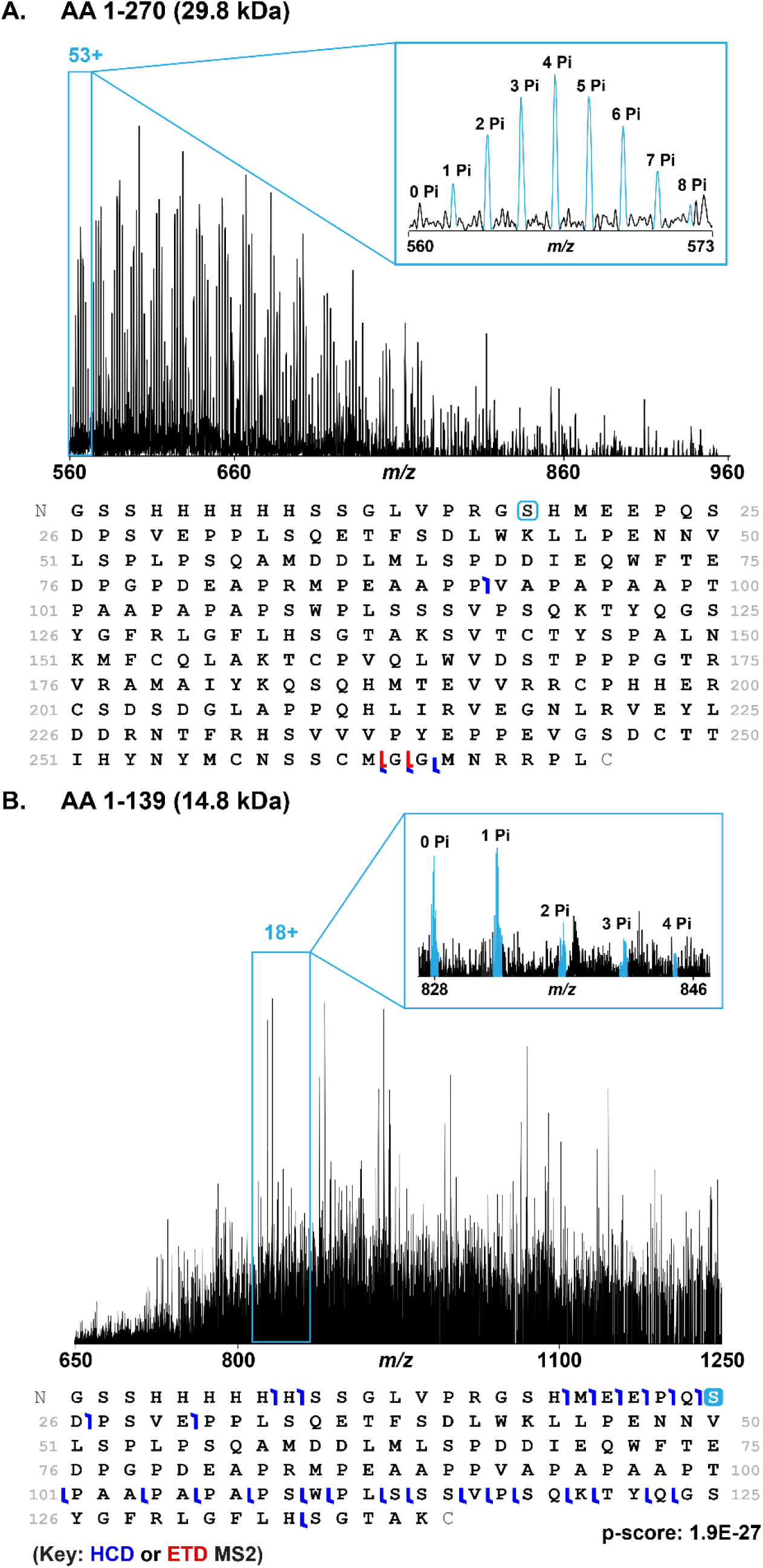
Visualizing phosphorylated sites within N-terminal regions of rp53 by middle-down mass spectrometry. **A.** MS^1^ spectrum of the 29.8 kDa partial digestion product comprising the N-terminal 269 amino acid residues of Chk1 phosphorylated rp53 (top), with MS^2^ fragment ion map of the corresponding sequence and a hypothetical phosphorylation site (bottom). **Inset:** Zoomed-in view of the 53+ charge state, showing evidence for the presence of up to 8 phosphorylated sites. **B.** MS^1^ spectrum of the 14.8 kDa partial digestion product comprising the N-terminal 138 amino acid residues of Chk1 phosphorylated rp53 (top), with MS^2^ fragment ion map of the corresponding sequence and an example phosphorylation site (bottom). **Inset:** Zoomed-in view of the 18+ charge state, showing evidence for the presence of up to 4 phosphorylated sites.

**Figure S8.**
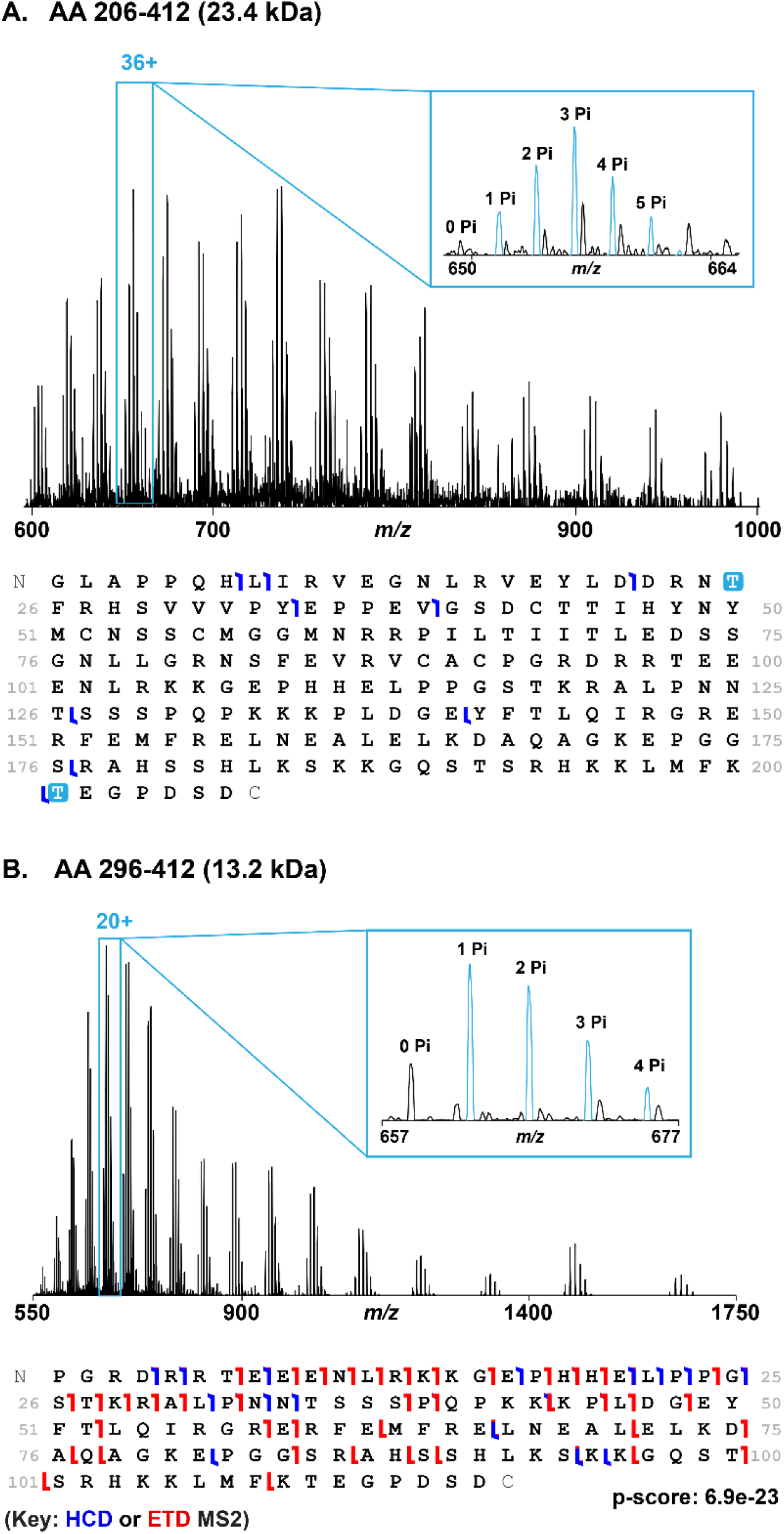
Visualizing phosphorylated sites within C-terminal regions of rp53 by middle-down mass spectrometry. **A.** MS^1^ spectrum of the 23.4 kDa partial digestion product comprising the C-terminal 206 amino acid residues of Chk1 phosphorylated rp53 (top), with MS^2^ fragment map showing partial characterization and example phosphorylation sites (bottom). **Inset:** Zoomed-in view of the 36+ charge state, showing evidence for the presence of up to 5 phosphorylated sites. **B.** MS^1^ spectrum of the 13.2 kDa partial digestion product comprising the C-terminal 116 amino acid residues of Chk1 phosphorylated rp53 (top), with MS^2^ fragment ion map of the corresponding sequence (bottom). **Inset:** Zoomed-in view of the 20+ charge state, showing evidence for the presence of up to 4 phosphorylated sites.

## Supplemental Tables

**Table S1.**
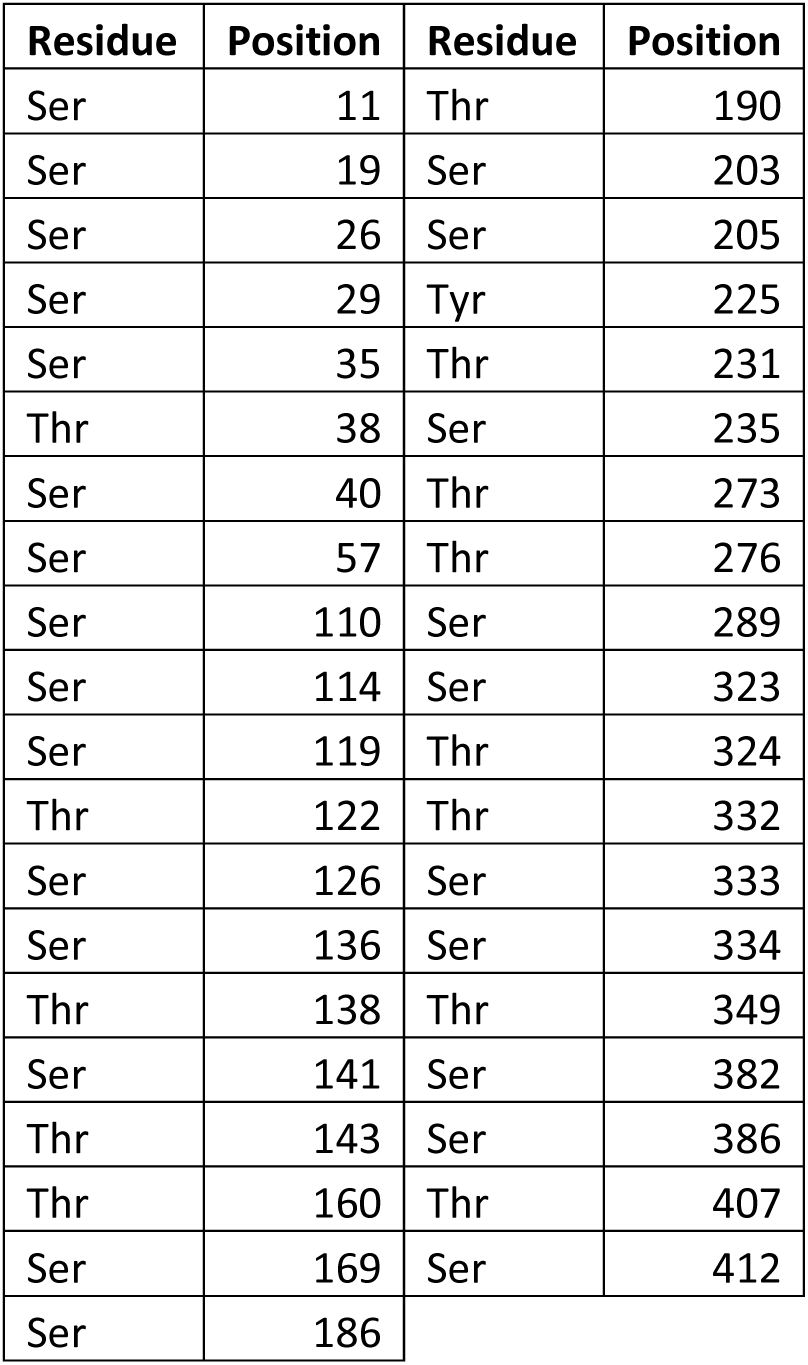
Phosphorylation sites mapped to rp53 by bottom-up mass spectrometry.

**Table S2.**
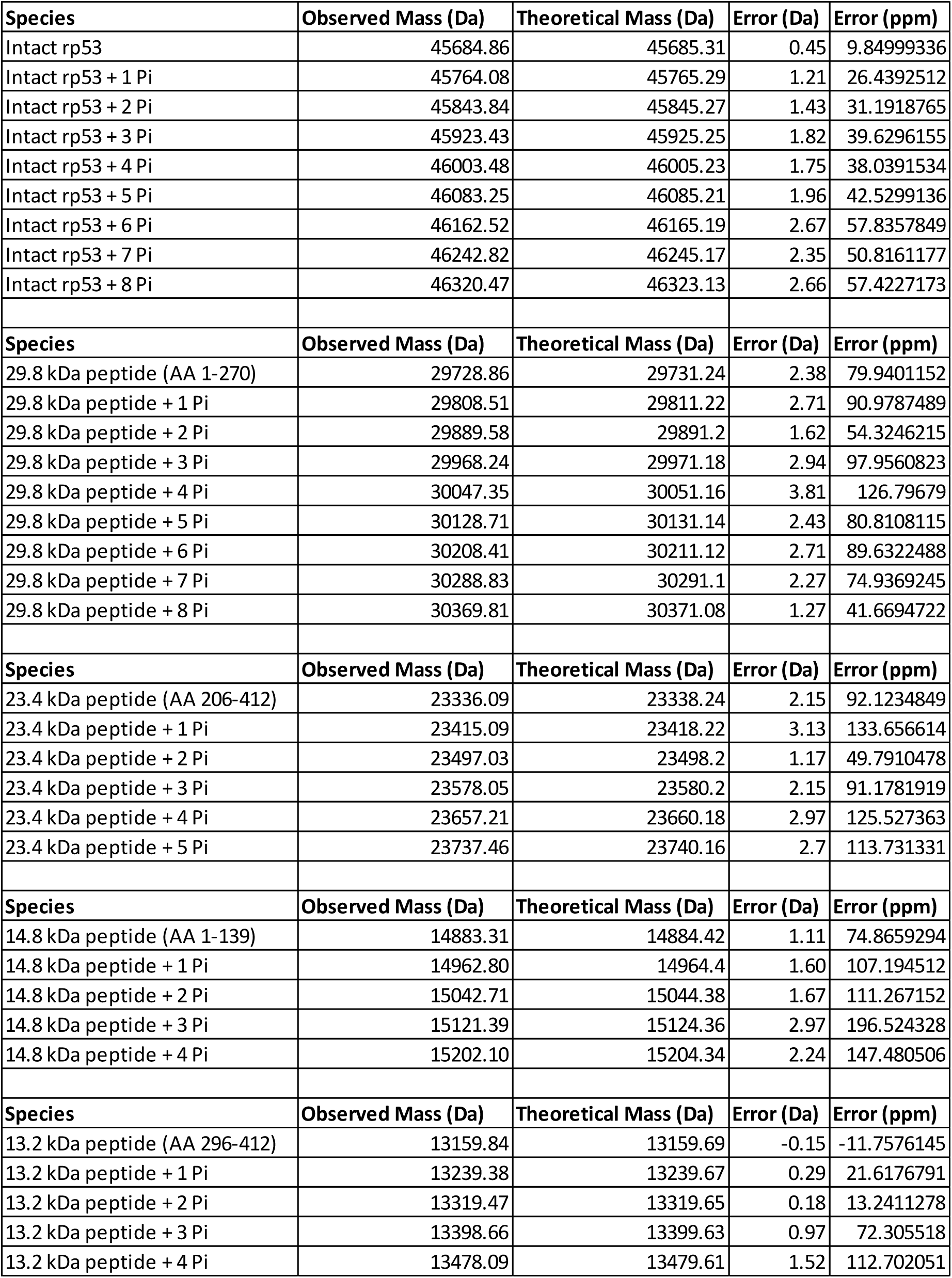
Deconvolution results from TD and MD MS of phospho-rp53 modforms.

**Table S3.** Estimation of phospho-rp53 modform distributions.

| Relative Intact rp53<br>Modform<br>Abundance (%) | Number of<br>Modification<br>Sites (N) | Number of<br>Potential<br>Modforms<br>(2<sup>N</sup>) | Estimated<br>Modform<br>Distribution<br>(%)*(2<sup>N</sup>) |
| --- | --- | --- | --- |
| 0.03 | 0 | 0 | 0 |
| 0.45 | 1 | 2 | 1 |
| 1.00 | 2 | 4 | 4 |
| 0.93 | 3 | 8 | 7 |
| 0.50 | 4 | 16 | 8 |
| 0.32 | 5 | 32 | 10 |
| 0.18 | 6 | 64 | 12 |
| 0.12 | 7 | 128 | 15 |
| 0.04 | 8 | 256 | 10 |
|  |  | **Total:** | **68** |
| Relative 29.8 kDa<br>Peptide Modform<br>Abundance (%) | Number of<br>Modification<br>Sites (N) | Number of<br>Potential<br>Modforms<br>(2<sup>N</sup>) | Estimated<br>Modform<br>Distribution<br>(%)*(2<sup>N</sup>) |
| 0.06 | 0 | 0 | 0 |
| 0.64 | 1 | 2 | 1 |
| 0.59 | 2 | 4 | 2 |
| 1.00 | 3 | 8 | 8 |
| 0.67 | 4 | 16 | 11 |
| 0.94 | 5 | 32 | 30 |
| 0.34 | 6 | 64 | 22 |
| 0.26 | 7 | 128 | 33 |
| 0.05 | 8 | 256 | 13 |
|  |  | **Total:** | **120** |
| Relative 23.4 kDa<br>Peptide Modform<br>Abundance (%) | Number of<br>Modification<br>Sites (N) | Number of<br>Potential<br>Modforms<br>(2<sup>N</sup>) | Estimated<br>Modform<br>Distribution<br>(%)*(2<sup>N</sup>) |
| 0.04 | 0 | 0 | 0 |
| 0.21 | 1 | 2 | 0 |
| 0.63 | 2 | 4 | 3 |
| 1 | 3 | 8 | 8 |
| 0.6 | 4 | 16 | 10 |
| 0.15 | 5 | 32 | 5 |
|  |  | **Total:** | **25** |

